# Time restricted feeding ameliorates colitis through clock-dependent and nutrition- dependent processes

**DOI:** 10.64898/2026.08.28.747868

**Authors:** V. Carmona-Alcocer, T. Igbokwe, J. MacDonald, M. Yousif, A. Todorovski, J. Grondin, W. Khan, P. Karpowicz

**Affiliations:** Department of Biomedical Sciences, University of Windsor, Windsor, ON, Canada; Department of Biology, Washington University, St. Louis, MO, USA; Department of Pathology and Molecular Medicine, McMaster University, Hamilton, ON, Canada; Farncombe Family Digestive Health Research Institute, McMaster University, Hamilton, ON, Canada

## Abstract

Inflammatory Bowel Disease (IBD) is rising worldwide and requires the development of preventative strategies to lower its incidence. Circadian rhythms, daily cycles of physiology, have been shown to be implicated in IBD in both clinical and laboratory studies. We tested if a 12-hour Time-Restricted Feeding (TRF) dietary regimen, that improves circadian rhythms, could ameliorate colitis using a mouse model of disease. We find that TRF indeed protects mice from the damaging effects of colitis, and boosts the rhythmic expression of transcripts related to the clock, cell proliferation, and barrier function. Under TRF, mice exhibit lower disease symptoms, changes in microbiota, and reduced inflammatory tissue damage. These outcomes are not clock-dependent: colitis in *Bmal1* mutants without circadian rhythms is also rescued by TRF. However, in this context alterations to the transcriptome are distinct from that found in clock-wildtype controls. Together our data support the translation of a 12-hour TRF in IBD, and highlight clock-dependent and nutrition-dependent changes accompanying colitis rescue.

## Introduction

Inflammatory Bowel Disease (IBD) is a chronic illness of the gastrointestinal tract whose incidence is increasing, particularly in developed countries where it is projected to reach >1% of the total population^1,2^. A form of IBD, Ulcerative Colitis, affects the colon, a complex organ that is home to an abundant microbiome. Colitis is thought to arise from complex interactions of environmental and genetic factors that shape human responses to microbiota, and whose dysfunction leads to inflammatory events^3^. A pressing need for preventative measures to lower IBD development, or to treat existing IBD, is required^1^. In addition, understanding the factors that promote IBD etiology is critical to explain how the incidence of this disease is rising.

Circadian rhythms are daily oscillations in physiology that include sleep/wake, body temperature, hormone, and feeding / fasting changes^4,5^. These are controlled by a conserved molecular system of transcriptional and post-transcriptional regulatory processes called the circadian clock. The clock consists of the transcription factors *Clock* and *Bmal1* that drive the expression of their negative regulators *Per1-2* and *Cry1-2*, as well as *Nr1d1-2* and *Ror* (family) factors, to generate 24-hour rhythms in transcription / translation^4^. Circadian clock rhythms in tissue cells ensures that these are synchronized with respect to the external daily environment, and in relation to one another throughout the body. Circadian rhythms promote health in the gastrointestinal tract^5,6^ and their disruption elevates illnesses including IBD^7,8^. IBD severity is higher in shift-workers who undergo circadian-disruption^9–11^, and IBD patients express aberrant levels of clock genes^9,12,13^. Indeed, disruptions in circadian cardiac rhythms occur prior to inflammatory flare-ups in IBD patients, and can thus predict disease onset when patients wear health monitoring devices^14^. Consistent with these clinical reports, studies using mouse models of colitis have confirmed that circadian clock mutants have increased disease^15–18^. The circadian clock has emerged as an important factor in IBD, whose activity could be harnessed to ameliorate disease outcomes.

Time-Restricted Feeding (TRF) is the synchronization of feeding / fasting cycles together with circadian physiological rhythms^19^. In mouse models, TRF evokes a cellular response that enhances clock strength and coordinates the daily timing of liver transcription, thereby rescuing metabolic physiology when mice are fed high fat diets^20–22^. The intestine is particularly responsive to the time of feeding, which induces shifts in the rhythms of genes including clock components^23,24^ and non-clock components^25^. It is not clear to what extent circadian versus non-circadian mechanisms underlie the transcriptional changes arising from timed feeding / fasting. Nonetheless, TRF has been found to exert a potent effect on mouse models of IBD: lowering inflammation, preserving tissue integrity, and improving recovery^26–29^. These pre-clinical TRF mouse studies showed that a 4 to 8 hour feeding window (corresponding to a 20 to 16 hour fasting period) is protective in the intestine. Indeed, an 8-hour restricted eating pilot study conducted for 30 days showed anti-inflammatory and microbial effects in human subjects^30^. However, normal human 24- hour eating habits are variable, with an individual’s eating intervals (first to last meal) ranging from 11-16 hours in duration^31^. A significantly shorter 4 to 8 hour feeding window is not likely to be palatable as a general long-term anti-IBD strategy.

To address this problem, we performed a study using mice to determine whether a 12- hour TRF regimen improves colitis outcomes. This window of time spanned from the early morning (*i.e*. 7am) to evening (*i.e*. 7pm), an interval that is sustainable for IBD patients. We found that 12-hour TRF greatly reduces colitis in wildtype mice, strengthening circadian clock gene expression, boosting the daily rhythms of >1700 genes that include cell cycle regulators and immune system components, and eliciting beneficial changes in the microbiome. We further tested if these effects are clock-dependent, using the *Bmal1* null mutant mouse strain. We predicted that the circadian system is needed mechanistically to drive TRF-induced transcriptional rhythms. Unexpectedly, *Bmal1* mutants display the same benefits of TRF, indicating that the circadian clock is not required. However, the transcriptional outcomes in this clock-disrupted context are distinct, with a completely different genetic program and calorie restriction phenotype accompanying the colitis-rescuing effects of TRF. Our findings support the use of a 12- hour eating / fasting strategy in lowering colitis in both circadian-competent and circadian-disrupted (shift worker) contexts.

## Results

### Time-Restricted Feeding (TRF) protects against colitis

We tested whether a 12-hour TRF regimen, that limits food access to an individual’s active phase, affects IBD disease susceptibility using a mouse model of colitis^32^. Wildtype mice (*Bmal1+/+*) were maintained under *ad libitum* (AL) or TRF, 2 weeks prior to colitis induction, then tested for disease symptoms, behaviour, and tissue analysis (Fig 1a). All mice were maintained under 12-hour light / 12-hour dark photoperiod, and TRF was provided from ZT12 to ZT24 (active phase of nocturnal mice, ZT is Zeitgeber Time which is respective to lights on at ZT0 and lights off at ZT12). TRF significantly ameliorates colitis severity, as indicated by a lower disease score (includes weight loss, bleeding, diarrhea, *etc*.) compared with AL animals (Fig 1b, ySupplementary Fig 1a). For instance, TRF reduces the number of individuals with blood in the stool at day 4, by a full order of magnitude, and rectal bleeding at day 6, by an even higher amount (Fig 1c). This is not due to the absence of damage in the colon during colitis, as both conditions cause a reduction in colon length due to inflammation (Supplementary Fig 1b). However, histological analysis indicates the TRF colon shows fewer ulcerated lesions with amorphous structure in the distal colon (Fig 1d-e), consistent with the lower disease symptoms apparent. Along with the reduced lesions, crypt length is protected in mice under TRF (Supplementary Fig 1c-g), indicating improved maintenance of epithelial structure during colitis. Together, these data show that a 12-hour TRF reduces colitis and preserves colon tissue architecture.

**Figure 1.**
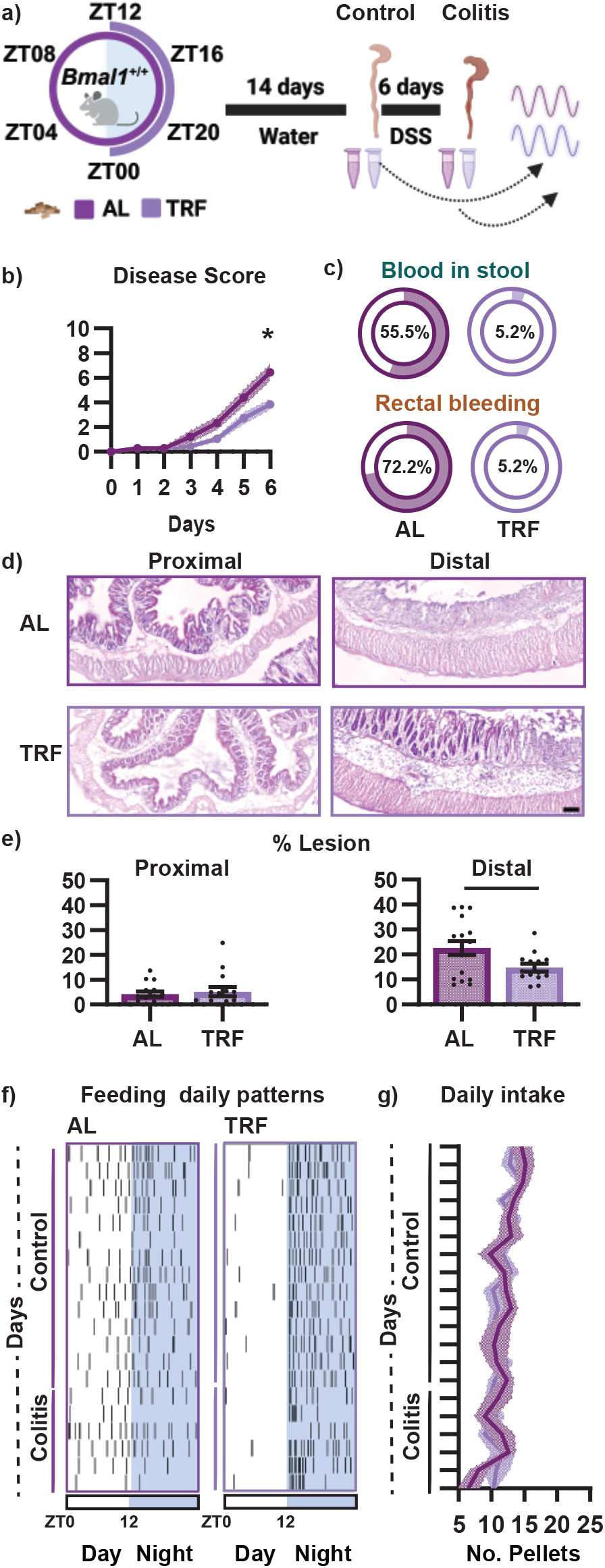
Time-Restricted Feeding (TRF) protects against colitis. **a)** Schematic showing experimental design. Mice with a functional clock (*Bmal1+/+*) were assigned to either *ad libitum* feeding (AL, purple) or time-restricted feeding (TRF, violet). Control (undamaged) and Colitis conditions were tested under AL *vs*. TRF. **b)** *Bmal1+/+* mice under TRF exhibited lower disease activity index compared to AL animals (day 6, p = 0.0163, two-way ANOVA, Sidak’s multiple comparisons test). **c)** For example, TRF reduces the number of individuals with blood in stool (day 4, p=0.0026) and rectal bleeding (day 6, p<0.0001, t-test). **d-e)** TRF decreases epithelial injury in the distal colon. **d)** Representative H&E from proximal colon (left) and distal colon (right) under AL (top) and TRF (bottom) conditions. **e)** Percentage of lesioned areas shows a significant decrease of in the distal colon under TRF (p=0.0267, unpaired t-test). **f-g)** TRF does not change daily feeding in *Bmal1+/+* WT mice. **f)** Representative daily feeding recordings for AL (left) and TRF (right). X-axis indicates 24-hour time, ZT0 (onset of the light), ZT12 (offset of the light); Y-axis days. **g)** No differences in total food intake between AL and TRF groups (two-way ANOVA). Data show mean ± s.e.m., n=18 per group.

We asked whether daily rhythms in food intake and locomotor behaviour are affected under a 12-hour TRF regimen. Although TRF alters the timing of food intake during the day, total caloric intake is equal between AL and TRF conditions (Fig 1f-g, Supplementary Fig 1h). AL mice show a characteristic increased food intake in the early evening just before lights are turned off (∼ZT12) and eat frequently during the day; TRF mice show a strong peak of feeding at ZT12 after having fasted during the whole day (Supplementary Fig 1i). During colitis these temporal patterns of feeding remain stable, suggesting that colitis itself does not disrupt daily feeding rhythms in either AL or TRF condition. Analysis of locomotor behavior, a measure of circadian rhythmicity, revealed that no significant differences between AL and TRF mice under control or colitis conditions are present (Supplementary Fig 1bj-k). Together, these results demonstrate that 12-hour TRF reduces colitis independently of changes in total food consumption, with rhythms in feeding and behaviour intact.

### TRF protects clock gene rhythms, during colitis, and increases daily gene expression rhythms

We, and others, have found that colitis dampens daily clock gene expression rhythms^16,18^, therefore we examined the 24-hour gene expression landscape in the colon under AL and TRF conditions. Distal colon tissue was collected over 24-hours (ZT0, 4, 8, 12, 16, 20), analyzed by RNA-sequencing, and diurnal rhythmicity was assessed using JTKCYCLE^33^. First, we asked whether TRF could prevent the dampening of clock gene oscillations during colitis (Fig 2a). Clock genes were analyzed using circular plots, where the time of maximum (peak) expression is represented over 24-hours and the distance from the centre is the amplitude of the cosinor fitted rhythm between maxima and minima. Under control conditions, prior to the induction of colitis, both AL and TRF animals shown robust daily oscillations in clock genes (Fig 2b). Next, we compared the timing of clock gene rhythms, which have a characteristic structure as positive and negative regulators of transcription drive gene expression^4^. In control conditions, *Bmal1* peaks in the morning, *Nr1d1* later in the day, *Per1-2* peaks in the early evening followed by *Cry1*. The timing of these genes is highly similar between AL and TRF mice, and is consistent with previous RNA-seq analyses of the intestine or colon, indicating a normal circadian oscillation^28,34–37^. This structure is dampened or lost during colitis: the AL group loses circadian clock gene rhythms in all genes except *Bmal1*, *Per2-3*, and *Nr1d2*. However, TRF is able to restore 24-hour oscillation in *Npas2*, *Nr1d1*, *Per1*, *Cry1*, and *Rorc* (Fig 2b-c, Supplementary Fig 2), indicating a boost to clock expression rhythmicity. These results indicate that while colitis reduces the number of rhythmic clock genes under AL feeding, TRF protects the molecular clock.

**Figure 2.**
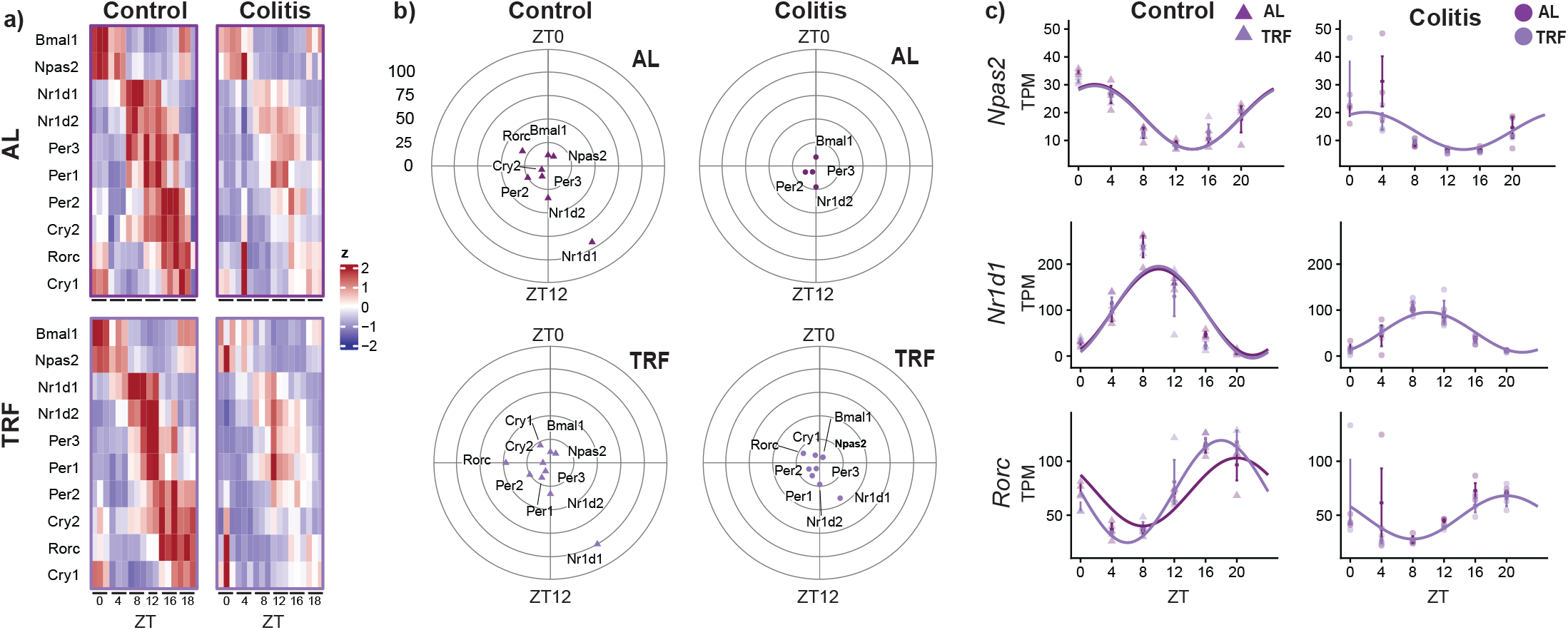
TRF protects 24-hour clock gene rhythms during colitis. **a)** Heatmaps display clock gene expression across 24-hours (Z-score scaled, n=3 individuals for each time point). Core clock genes in control and colitis conditions under AL or TRF show daily changes that are overall reduced during colitis. **b)** Circular plots show circadian clock genes identified as rhythmic by JTK Cycle (AL are triangles, TRF are circles). Phase, shown as peak time of the 24-hour rhythm, is mapped over the day (ZT0–ZT24), and rhythm amplitude is the radial distance from the centre. More rhythmic clock genes are detected under TRF (lower) compared to AL conditions (upper). **c)** Graphs show examples of clock genes (*Npas2*, *Nr1d1*, and *Rorc*), that lose 24-hour rhythm during colitis, but are protected by TRF to remain in the same phase (timing). All graphs show TPM mean ± SEM values across the day. Rhythmic expression was assessed using JTK_CYCLE, and cosine fits are overlaid for genes classified as rhythmic (BH.Q < 0.1, amplitude > 0.05).

Next, we asked if TRF has a broader effect in driving rhythms in the colonic transcriptome. JTKCYCLE was again used to identify all 24-hour rhythmic genes in the distal colon tissue in samples tested by RNA-sequencing. Under control conditions, AL fed mice express genes throughout the day with >100 genes identified as rhythmic, TRF substantially increases the number of rhythmic transcripts to >1700 genes (Fig 3a). TRF also consolidates gene expression rhythms into a biphasic distribution that peaks at midday (ZT6-8) and early morning (ZT22-0/24). These changes are the result of many rhythmic genes peaking at these times. For instance, we detected that at midday the AL colon has >35 rhythmic genes with maxima at ZT6-8, but the TRF colon has >550 genes at the same time. This indicates that restricting food intake to the active phase imposes temporal gene expression changes in the colon. Similar to its dampening of circadian clock gene rhythms, colitis greatly reduces rhythmic gene expression under both feeding schedules. In AL colitis, the colon expresses very few rhythmic genes (11 identified), and under TRF colitis this number is increased marginally (46 identified). We also noted that the timing of rhythms is altered during inflammation, with peaks at the evening (ZT12-16) in both AL and TRF, and a second peak of rhythmic expression in the early morning (ZT0-2) in TRF conditions only (Fig 3a). Together this analysis demonstrates that TRF increases rhythmic gene expression, while colitis suppresses it.

**Figure 3.**
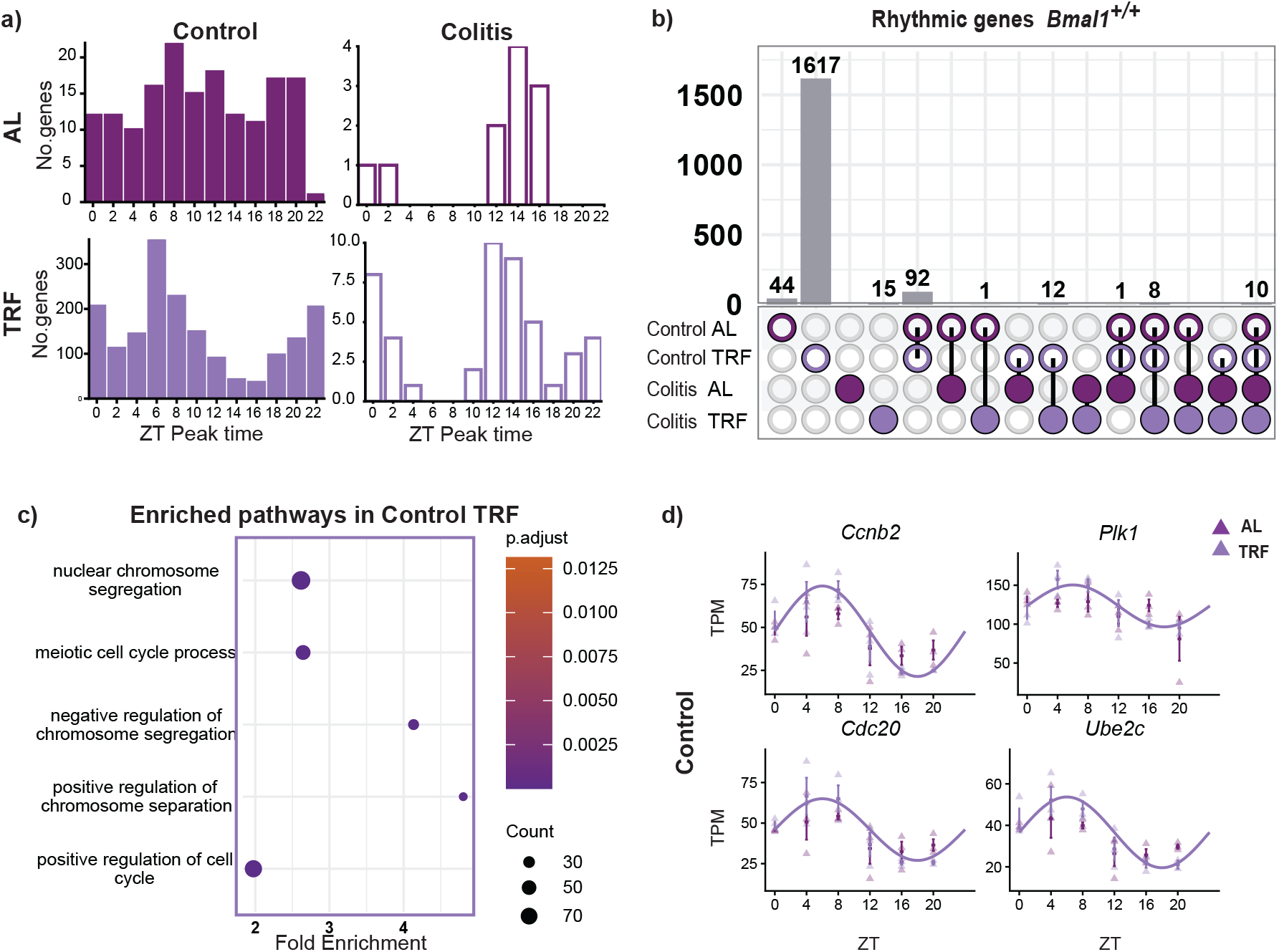
TRF increases daily gene expression rhythms. **a)** The TRF-treated colon shows a higher number of 24-hour rhythmic genes. Histograms show the peak of genes in the four conditions tested (X-axis indicates time of maximum expression by cosinor fit). Under control conditions, TRF shows many more rhythmic genes (compare the Y-axis in upper and lower graphs). Fewer genes are rhythmic during Colitis, with TRF showing a higher number. **b)** Upset plot showing number of rhythmic genes unique to each condition *vs*. shared. Few rhythmic genes are shared between AL and TRF, suggesting TRF induces rhythms in specific genes. **c)** Dot plot from ORA enrichment analysis showing top 5 GO terms: daily rhythms in cell cycle pathways are elevated in undamaged control animals under TRF. Dot size indicates number of genes, X-axis shows fold enrichment, and color the adjusted P value. **d)** Cell cycle genes that display daily rhythms under control TRF peak together in phase during the afternoon (∼ZT6). These include genes involved in mitotic progression (*Ccnb2*, *Plk1*, *Cdc20*, *Ube2C*). Graphs show TPM mean ± SEM values across the day. Rhythmic expression was assessed using JTK_CYCLE, and cosine fits are overlaid for genes classified as rhythmic (BH.Q < 0.1, amplitude > 0.05).

Are the daily rhythms in different conditions due to the same transcripts? To address this question, unique *vs*. shared genes were compared across all three conditions (Fig 3b, Supplementary Table 2). Very few rhythmic genes are common, indeed, 1617 rhythmic genes in TRF control conditions are unique with no overlap in any of the other conditions tested. 92 genes were shared in both control AL and TRF, 12 genes are shared by control and colitis TRF, and no genes were shared in colitis AL and TRF. Indeed the number of rhythmic genes detected in colitis conditions is relatively low, the 10 transcripts that are shared by these and both the control conditions (all 4 groups) are circadian clock genes. These findings suggest that TRF can drive temporal gene rhythms in the colon as a distinct rhythmic transcriptional program.

Given the large number of genes that show daily oscillation under TRF control conditions we performed Over-Representation Analysis (ORA) to detect enriched pathways in this data set (Fig 3c, Supplementary Table 2). We predicted that rhythmicity in TRF-induced rhythmic genes is protective prior to the induction of colitis. Cell division related pathways emerged as the 5 most significantly enriched functional categories, suggesting that TRF promotes the temporal coordination of proliferation in the undamaged control colon. Genes involved in cell cycle progression and mitotic regulation exhibit 24-hour rhythms in TRF-fed animals, these include *Ccnb2* and *Plk1* (G2/M phase progression), as well as *Cdc20* and *Ube2c* (APC/C-dependent mitotic progression) (Fig 3d). Of note, these transcripts of mitotic regulators are all expressed in the same phase, peaking together at ZT4-8 in the middle of the day, and none of these show rhythmicity during colitis in either AL or TRF (Supplementary Fig 3a). We have previously noted the rhythmic timing of cell proliferation in the colon^18^, and these data suggest that TRF may exert a pre-colitis protective effect through cellular regeneration.

We next examined the 12 genes that maintain rhythmic expression across both control and colitis TRF (Fig 3b, Supplementary Fig 3b, Supplementary Table 2). These include genes previously implicated in colon inflammation such as *Dusp6* (a MAPK pathway repressor^38,39^) and *Egln3* (a NFκB pathway repressor^40,41^), and in immune cell function such as *Fcho1* (a regulator of T-cell endocytosis^42,43^). This suggests TRF promotes rhythms in cell signaling and inflammatory functions. In addition to preserving gene rhythms, TRF induces *de novo* rhythmic oscillation in another 15 genes during colitis that include *Ank3* (a regulator of the intestinal barrier^44^) and *Gcnt3* (a regulator of mucus glycosylation^45^) (Fig 3b, Supplementary Fig 3b, Supplementary Table 2). Together, these results show that TRF induces rhythms in gene expression linked to colon physiology.

### TRF attenuates colitis in a clock-independent manner

Our investigation of clock wildtype (*Bmal1+/+*) mice indicates that TRF protects against colitis (Fig 1), and increases the daily rhythms of both circadian and non-circadian genes (Fig 2-3). We therefore asked if an intact molecular clock is required for the protective effects of TRF, essentially, if colitis protection arises from imposing non-clock rhythms on gene expression. To test this, *Bmal1-/-* null mutant mice, that have no transcriptional circadian rhythms^46^, were subjected to the same experimental feeding paradigm and tested upon the induction of colitis (Fig. 4a). Unexpectedly, TRF reduces disease severity in *Bmal1-/-* mice, these exhibit lower disease scores (Fig 4b) like the *Bmal1+/+* wildtype mice. Colon tissue analysis further confirmed that TRF attenuates colitis in the absence of a functional clock. For instance, colitis reduces colon length in AL *Bmal1-/-* mice, whereas TRF prevents this shortening (Fig 4c). Similar to wildtype *Bmal1+/+* mice (Fig 1d-e), TRF also reduces the frequency of lesions in the distal colon of *Bmal1-/-* mutants (Fig 4d-e, Supplementary Fig 4a). Epithelial structure was also maintained in *Bmal1-/-* TRF: during colitis, AL Bmal*1-/-* mice have reduced distal crypts, whereas TRF preserves distal crypt structure (Supplementary Fig 4b-e). These results mirror the phenotypes observed in wildtype *Bmal1+/+* mice and support the protective effect of TRF during colitis in *Bmal1-/-* clock-disrupted mutants.

**Figure 4.**
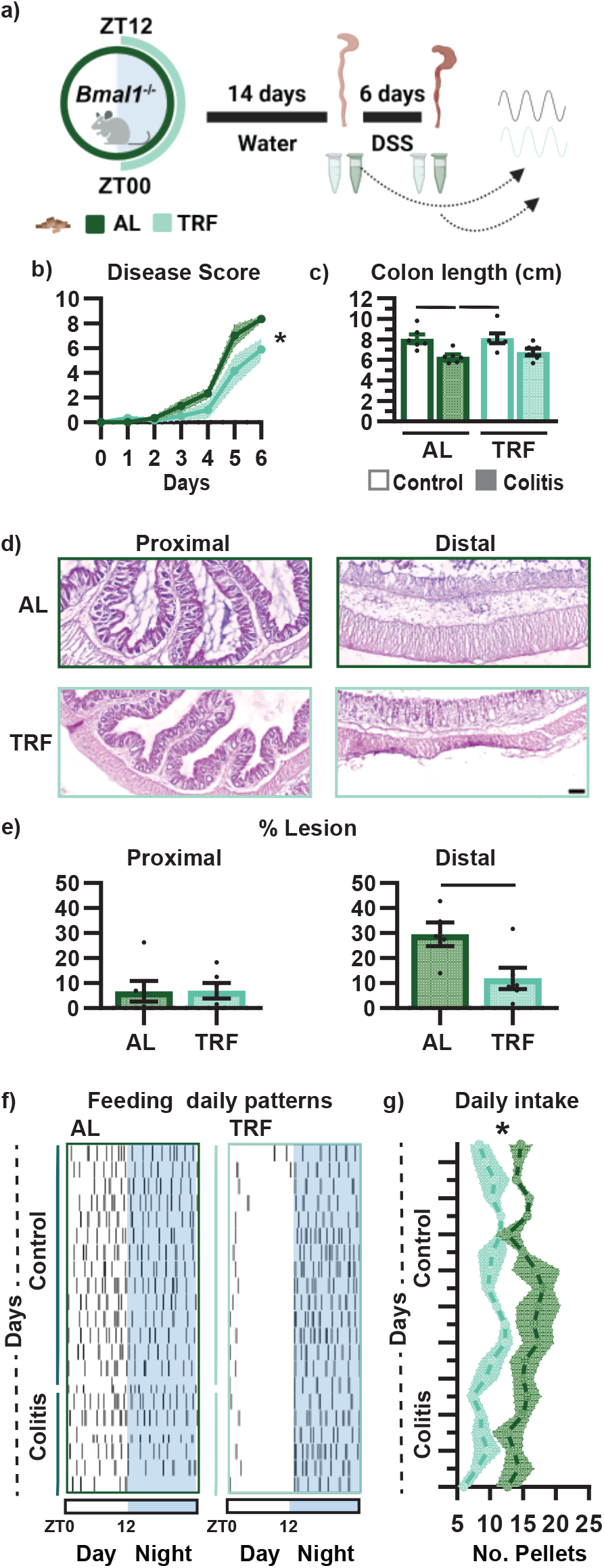
TRF attenuates colitis in a clock-independent manner. **a)** Schematic showing experimental design. Mice lacking a functional clock (*Bmal1-/-*) were maintained on AL (dark green) or TRF (light green), and then tested in Control (undamaged) *vs*. Colitis conditions. **b)** *Bmal1-/-* TRF mice exhibit lower disease compared to AL in colitis (p=0.0284, two-way ANOVA repeated measures), demonstrating the clock is not required for the beneficial effects of TRF in colitis. **c)** TRF prevents colon shortening during colitis in *Bmal1-/-* mice. Bmal1-/- mice have a significant decrease in colon length during colitis (p=0.0184, two-way ANOVA Tukey’s comparison test) that is prevented by TRF (no significant difference). **d-e)** TRF decreases epithelium damage in the distal colon of *Bmal1-/-* mice. **d)** Representative H&E of proximal and distal colon. **e)** Percentage of lesion area shows significant reduction in lesions in the distal colon under TRF (p=0.0303, Mann-Whitney test). **f-g)** In *Bmal1-/-* mice, imposing a TRF pattern prevents feeding during the day which causes caloric restriction. **f)** Representative daily feeding recordings for AL (left) and TRF (right); X-axis indicates 24-hour time, Y-axis days. *Bmal1- /-* mice eat throughout 24-hours, daytime feeding is prevented by TRF. **g)** *Bmal1-/-* have a decrease in total food intake under TRF (p=0.0065, two-way ANOVA repeated measurements, n=3 by group). Unless otherwise indicated, data show mean ± s.e.m., n=6 by group.

Due to the absence of circadian clock transcription, *Bmal1-/-* mice lack circadian rhythms in physiology and behaviour^46^. We tested feeding and locomotor activity to determine whether these were affected by TRF, reasoning that if TRF imposed daily rhythms in behaviour these could potentially accounting for the colitis protective effects. Under AL conditions, *Bmal1-/-* eat continuously, with TRF restricting food access during the day (Fig 4f). Unlike *Bmal1+/+* mice, which maintain equal caloric intake under AL and TRF conditions (Supplementary Fig 1h), *Bmal1-/-* mice decrease their intake when subjected to TRF (Fig 4g, Supplementary Fig 4f). This reduction in caloric intake is clear when examining average *Bmal1-/-* feeding patterns over 24-hours in AL *vs*. TRF conditions. AL *Bmal1-/-* mice eat uniformly throughout the day and night, under TRF daytime feeding is abolished, and food consumption patterns are similar before and during colitis in each group (Supplementary Fig 4g). This confirmed that TRF imposes a 24-hour rhythm in feeding on *Bmal1-/-* mice and revealed that this results in reduced caloric intake. Similar to their arrythmic feeding under AL, *Bmal1-/-* locomotor activity lacks circadian organization (Supplementary Fig 4h-i). Of note, this is not affected by TRF, even though no food is consumed during the day, *Bmal1-/-* TRF locomotor activity remains uniform day and night over 24-hours. These results demonstrate that while TRF protects against colitis in a *Bmal1*-independent manner, it does not alter circadian locomotor behaviours. Instead, TRF imposes a limitation on daytime feeding, thereby reducing caloric restriction in circadian clock-deficient animals.

### TRF alters daily gene expression in a distinct manner, in clock functional versus clock disrupted conditions

We tested if distinct transcripts underlie the beneficial effects of TRF in a clock-disrupted context. Distal colon tissue was collected in the morning (ZT0) and evening (ZT12), then analyzed by RNA-sequencing to determine if post-feeding and post-fasting samples exhibit transcriptional changes in AL *vs*. TRF conditions. Many genes are upregulated in *Bmal1-/-* TRF mice compared to their AL counterparts, however, the AL colon exhibits a distinct profile (Fig 5a, Supplementary Table 3). ORA enrichment analysis indicated that *Bmal1-/-* AL colons have mostly genes related to ribosome function upregulated before colitis (Fig 5b). This is evident when chord plots showing the highest upregulated genes are linked to the ORA enriched pathways: many ribosome components (*Rpl* family genes) are highly expressed (Fig 5c). The *Bmal1-/-* TRF colon, on the other hand, is enriched in mitochondrial related genes such as *Cox2-3* (*Cytochrome Oxidase*) and *ATP6/8* (*ATP synthase*), or lipid transport such as *Fabp* (*Fatty Acid Binding Protein*) and *Mttp* (*Microsomal Triglyceride Transfer Protein*) (Fig 5d-e). Under colitis these alterations in gene expression are changed: the *Bmal1-/-* AL colon shows many upregulated genes that are absent in the TRF condition (Fig 5f). These genes include *Krt* (*Keratin*) family genes that function in differentiated epithelial barrier cells^47^, and *Cdk6* (*Cyclin Dependent Kinase 8*) that adapts cell growth to stress^48^ (Fig 5g-h); these are downregulated in the TRF condition, suggesting a lowered cellular differentiation state. Together, these data suggest the *Bmal1-/-* colon is reprogrammed by TRF to drive an aerobic metabolism program in lieu of cellular maturation.

**Figure 5.**
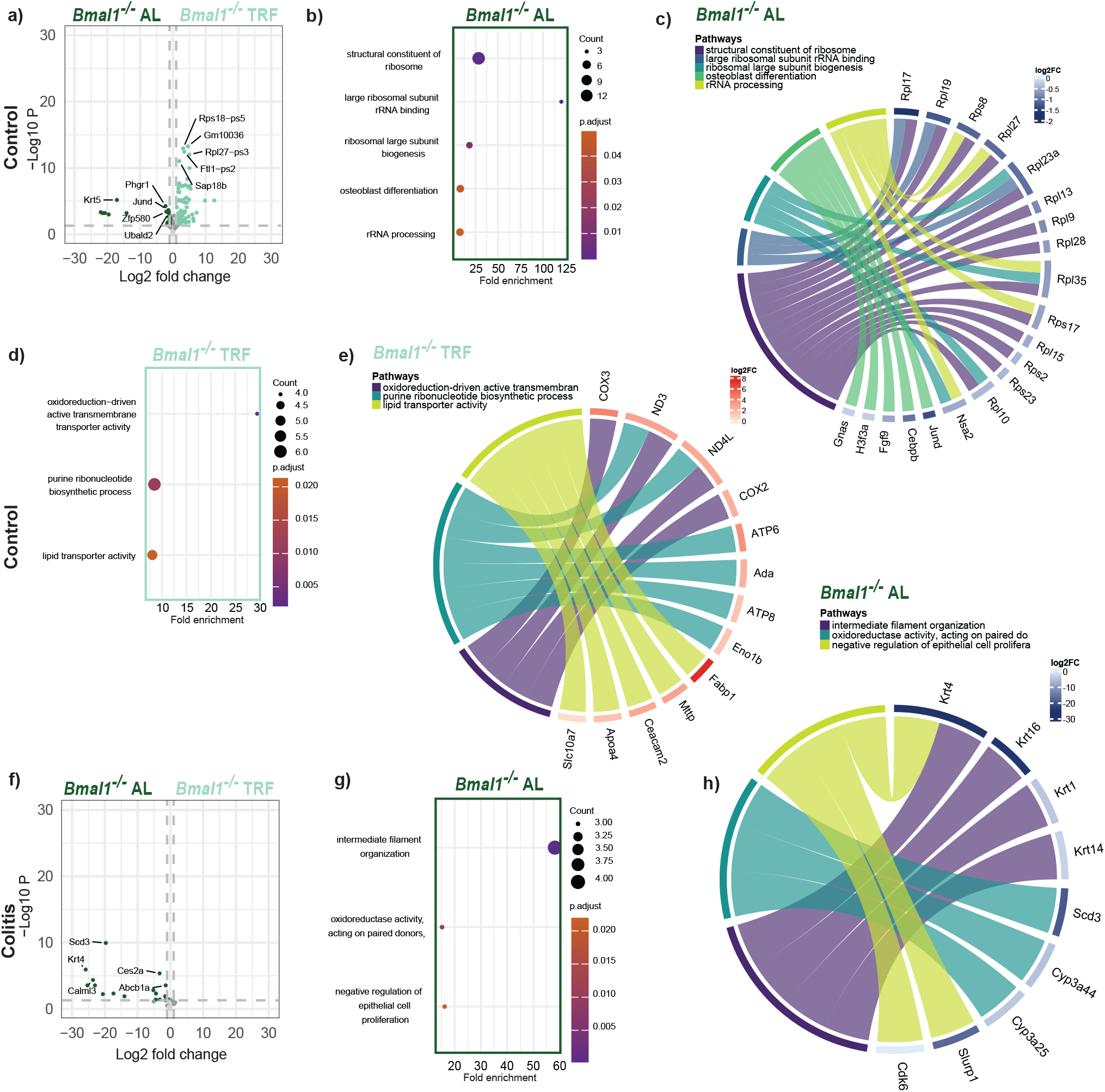
TRF alters gene expression in the *Bmal1-/-* colon to favour metabolic pathways. **a)** Differential gene expression analysis comparing the *Bmal1-/-* colon under AL *vs*. TRF (control conditions). **b)** Dot plot from ORA enrichment analysis showing top 5 GO pathways that are enriched in *Bmal1-/-* AL that include rhythms in ribosome- associated processes. **c)** Chord plot showing the connection between differentially expressed genes and these enriched pathways in *Bmal1-/-* AL. Chord colors correspond to pathways, and gene segments are colored to show log₂ fold change in expression. **d)** Dot plot from enrichment showing that *Bmal1-/-* TRF, unlike *Bmal1-/-* AL under control conditions, has rhythms in metabolic pathways (only 3 pathways detected). **e)** Chord plot showing examples of genes enriched in *Bmal1-/-* TRF, these include mitochondrial and lipid pathway genes. **f)** Differential gene expression analysis comparing the *Bmal1- /-* colon in AL *vs*. TRF during colitis. **d)** Dot plot from enrichment in *Bmal1-/-* AL colitis conditions reveal an increase in genes negatively regulating the cell cycle (only three pathways detected). No enriched pathways were detected in the *Bmal1-/-* TRF. **e)** Chord plot shows the connection between these differentially expressed genes and enriched pathways. Volcano plots show log₂ fold change versus –log₁₀ adjusted p-value, log₂ fold thresholds 0.6 applied. In dot plots, size represents gene count, and color indicates adjusted p-value.

We next revisited clock wildtype *Bmal1+/+* mice and compared these to clock-disrupted *Bmal1-/-* mice to see whether feeding and fasting induce clock-dependent changes in daily gene expression (Supplementary Fig 5). We compared differentially expressed genes in the undamaged (control) colon at ZT0 (after mice have been feeding all night) to ZT12 (after fasting, when food is first given to TRF mice). In the control state, the *Bmal1+/+* AL colon and TRF colon have differentially upregulated genes in the fed and fasted states, while the *Bmal1-/-* colon in both conditions only shows upregulated genes in the fasted state (Fig 6a). This suggests that transcriptional changes do not occur in these clock-disrupted mice after feeding. We performed ORA enrichment analysis (Supplementary Fig 6, Supplementary Table 4) and graphed chord plots for all four conditions to visualize the affected genes / pathways (Fig 6b-e). Under control conditions, *Bmal1+/+* AL mice express genes involved in cell division after feeding and circadian genes after fasting (Fig 6b). While DNA replication genes, related to cell division, are also upregulated post-feeding in *Bmal1+/+* TRF, these are joined by protein folding, heat shock, and GTPase related genes; *Bmal1+/+* TRF mice show histone modification, ion transport, and secretory genes upregulated post fasting (Fig 6c). These data reveal that in wildtype mice, the colon exhibits distinct gene expression profiles in fed / fasted states under TRF. In the *Bmal1-/-* mutant AL colon, ribosome related genes and mitochondrial / oxidative metabolic genes are upregulated in the fasted state (Fig 6d). The *Bmal1-/-* TRF colon, upregulates different genes after fasting, including *Krt* genes related to differentiation and genes related to immune system antibody responses (*Wfdc5*, *Spink5*, *etc*.) (Fig 6e). This indicates that TRF also creates distinct transcriptional programs in the clock-disrupted *Bmal1-/-* colon following a fasted state. Comparing genes irrespective of fed/fasted states supports the findings that unique transcriptional profiles are present in a clock-dependent manner. For instance, *Bmal1+/+* AL mice show more inflammatory activity than *Bmal1-/-* AL, and under TRF, *Bmal1+/+* upregulate metabolic genes (Supplementary Fig 5). Colon transcriptional profiles are distinct in the *Bmal1+/+* and *Bmal1-/-* genotypes.

**Figure 6.**
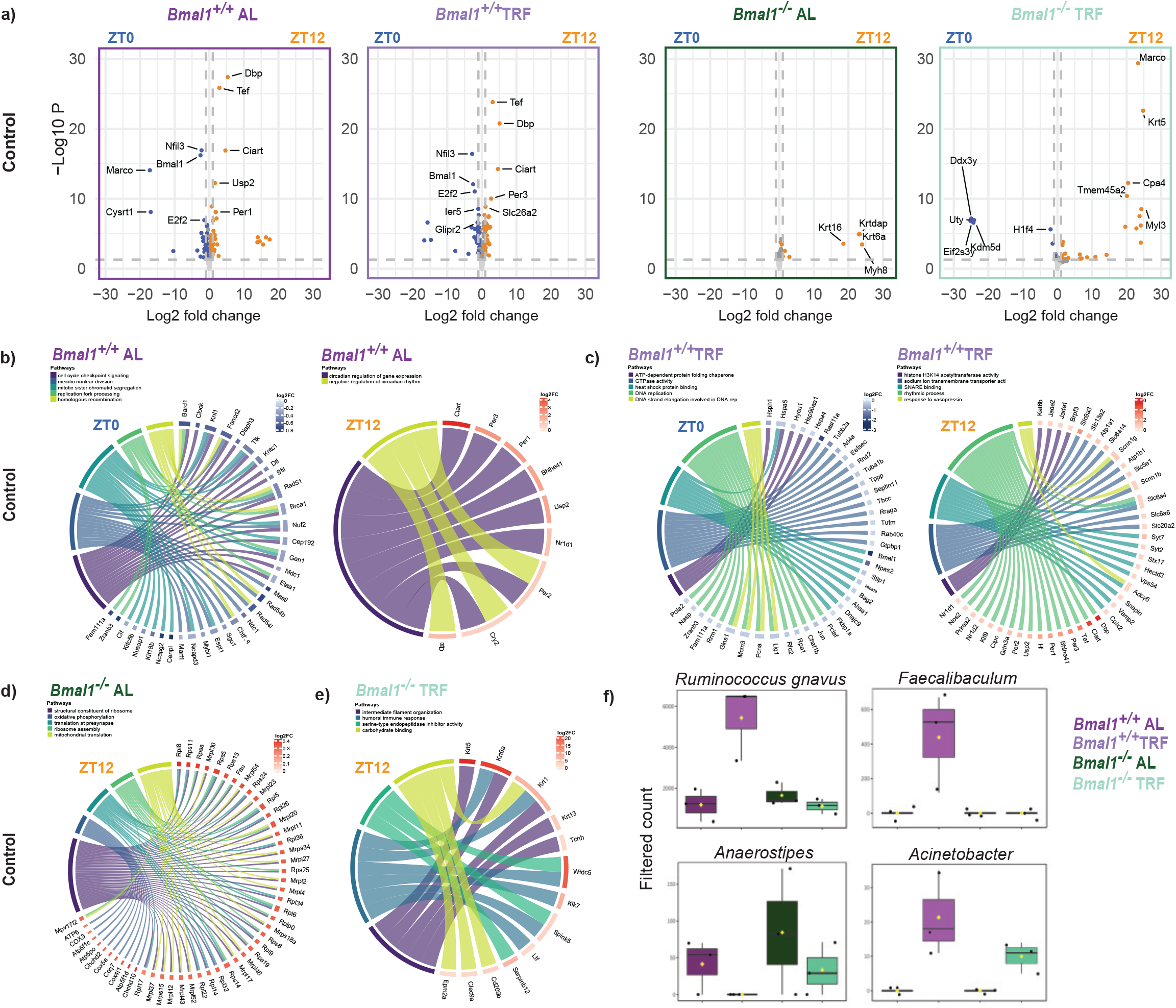
TRF reprograms feeding / fasting gene expression timing. **a)** Volcano plots show differentially expressed genes in ZT0 (blue: morning after mice have fed during the night) and ZT12 (orange: evening after TRF mice have fasted) in control conditions (no damage). The *Bmal1+/+* colon exhibits genes at both time points, note that at ZT0 in the *Bmal1-/-* colon there are very few genes differentially expressed. **b)** Chord plot shows examples of differentially expressed genes connected to enriched pathways. The *Bmal1+/+* AL colon shows enrichment of genes regulating cell division in the morning, and circadian genes in the evening (only two pathways detected). Chord colors correspond to pathways, and gene segments are coloured to represent log₂ fold change. **c)** Chord plot of same for the *Bmal1+/+* TRF colon that shows different pathways are enriched at these same times. The morning shows genes involved in protein production, as well as cell division, while the evening’s circadian gene program is joined by secretory (SNARE binding) and ion transport processes. **d)** Chord plot of enrichment in *Bmal1-/-* AL colon (ZT12 only) shows genes involved in ribosome function and aerobic metabolism. **e)** Chord plot for same in *Bmal1-/-* TRF colon (ZT12) shows that these processes are replaced by anti-inflammatory responses (*i.e.* humoral immune response) and differentiation (intermediate filament organization, *Krt* family genes). **f)** Microbial composition was assessed by 16s rRNA analysis and tested at ZT0 and ZT12. Certain bacteria associated with IBD are altered under TRF, including *Ruminococcus* and *Faecalibaculum* (higher in *Bmal1+/+* TRF), *Anaerostipes* (lower in *Bmal1+/+* TRF), and Acinetobacter (higher in both *Bmal1+/+* and *Bmal1-/-* TRF).

The colon is host to a large microbiome population, itself having daily rhythmicity, that is thought to be driven by food intake timing^49^. We therefore examined microbiome composition in both genotypes at ZT0 (fed) and ZT12 (fasted) using 16S rRNA sequence analysis (Supplementary Fig 7a). At the genus level, beta diversity measures indicated dissimilarities between *Bmal1+/+* AL compared to both *Bmal1+/+* TRF (p = 0.008, FDR= 0.016), *Bmal1-/-* AL (p=0.004, FDR =0.012), and *Bmal1-/-* TRF (p= 0.002 and FDR =0.012) conditions (Supplementary Fig 7b). No difference was found between any of the groups in terms of Alpha diversity via Shannon index (Supplementary Fig 7c). These changes are driven by alterations in specific bacterial populations. For instance, *Bmal1+/+* TRF have increased *Acinetobacter*, *Anaeroplasma*, *Parasutterella*, *Ruminococcus gnavus*, and *Faecalibaculum* (Fig 6f, Supplementary Tables 5-6). Of note, both *Ruminococcus* and *Parasutterella* have been positively associated with intestinal inflammation^50,51^. However, butyrate-producing *Faecalibaculum* has been shown to correlate with reduced colitis severity and improved barrier integrity in mouse models^52,53^. Further, in *Bmal1+/+* TRF mice, *Anaerostipes* was uniquely depleted compared to all other groups (Fig 6f, Supplementary Tables 5-6). *Anaerostipes* have been found to have a negative effect in colitis in both mice and humans^54,55^; depletion of this microbe may, in part, account for the decreased severity of colitis noted in this group. Of note, *Acinetobacter* was also higher in *Bmal1-/-* TRF mice, suggesting clock-disrupted mice also benefit from circadian food intake cycles. We further examined fed *vs*. fasted timepoints and noted that, upon fasting, both TRF genotypes showed discrete clustering in beta diversity with increased levels of *Anaeroplasma* and *Acinetobacter* as well as diminished levels of both *Lachnospiraeceae* and *Herbinix* upon fasting (summary and full statistical details are provided in Supplementary Tables 5-6). Overall, TRF imposes specific and shared microbiome changes in wildtype *Bmal1+/+* and clock-disrupted *Bmal1-/-* mice.

Do these changes persist during colitis? We repeated our transcriptional analyses from colon samples collected during colitis, to determine if these also show a TRF-dependent alteration. Differentially expressed genes are expressed at both fed and fasted states in *Bmal1+/+* AL and *Bmal1-/-* TRF mice, while *Bmal1+/+* TRF shows genes most differentially expressed only upon fasting (Fig 7a). *Bmal1-/-* AL has few genes changing at these times, likely reflecting its arrhythmic feeding pattern. ORA analysis (Supplementary Fig 8a-c) indicated that the *Bmal1+/+* AL and TRF colon expresses similar genes in coltis (Fig 7b-c) as it does in undamaged conditions (Fig 6b-c), however, in TRF the colon upregulates genes involved in xenobiotic pathways during colitis (Fig 7c). On the other hand, the *Bmal1-/-* TRF colon shows a curious pattern of upregulating colon tissue structure genes (extracellular matrix, adhesion, and vasculature) in the morning, following feeding at night, and cell division genes in the evening following fasting (Fig 7d). These data again show that TRF imposes a distinct timing on gene expression in this tissue, when *Bmal1* clock transcription is absent, that is not shared by the other conditions.

**Figure 7.**
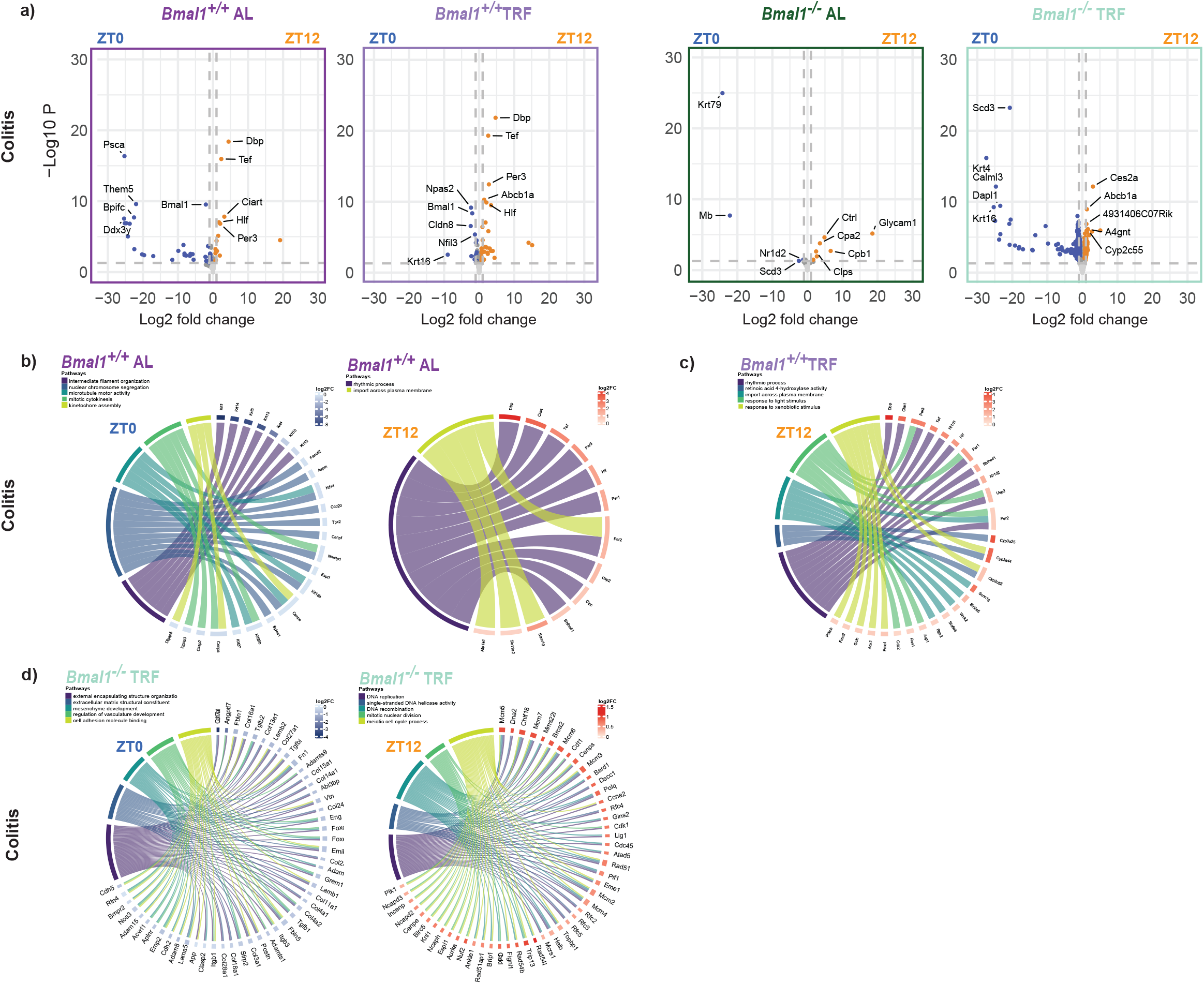
TRF imposes gene expression timing on the Bmal1-/- colon in colitis. **a)** Volcano plots show differentially expressed genes during colitis. As in Fig 6, comparisons were done between ZT0 (blue: morning post-feeding) and ZT12 (orange: post-fasting). Note that no comparisons could be made from the Bmal1-/- AL colon during colitis (very few differentially expressed genes), and only ZT12 could be analyzed in the Bmal1+/+ TRF colon (few genes detected at ZT0). **b)** Chord plot illustrating gene to enriched pathway associations for the *Bmal1+/+* AL colon during colitis, chord colors correspond to pathways, and gene segments are coloured to represent log₂ fold change. Similar to control (undamaged) conditions, the colon expresses cell division related genes at ZT0 and circadian genes at ZT12. **c)** Chord plot for ZT12 in the *Bmal1+/+* TRF colon during colitis (none detected at ZT0 in this condition). Similar pathways are detected as in control conditions (circadian and transport), but xenobiotic genes are also elevated (*i.e. Cyp* family). **d)** Chord plot for *Bmal1-/-* TRF colon during colitis. TRF reprograms the Bmal1- /- colon to express genes according to feeding / fasting times. At ZT0, genes in adhesion, stromal structure (extracellular matrix, and mesenchyme), and vasculature are upregulated, while at ZT12, genes in cell division are upregulated.

## Discussion

We find that a 12-hour TRF dietary regimen is sufficient to ameliorate colitis, greatly reducing disease symptoms and preserving colon tissue (Fig 1 and 4). These colitis-protective outcomes are clock-independent, as they occur in *Bmal1-/-* mutants that have no circadian clock gene expression rhythms. However, in wildtype *Bmal1+/+* mice, TRF protects circadian clock rhythms which are dampened during inflammation, and drives diurnal rhythms in hundreds of non-clock genes as well (Fig 2-3). Thus, TRF colitis rescue in *Bmal1+/+* mice may occur in part through the circadian transcriptional system. In support of this notion, we note that TRF-dependent genes are completely distinct in *Bmal1-/-* mutants (Fig 3, 5–7), suggesting that timed nutrition imposes a unique non-clock related transcriptional timing based on feeding responses. Our finding that in the wildtype colon, TRF enhances rhythms in cell division genes is consistent with previous studies showing rhythms in colon regeneration^18^ are enhanced by feeding times^56^. Of note, the *Bmal1-/-* mutant is enriched for cell division regulators in the early evening during colitis (ZT12, Fig 7d), an inversion of the enrichment observed in the wildtype (Fig 6b-c, 7b). This suggests the cell proliferation timing imposed by TRF organizes the colon but in an aberrant manner if the clock is not present. Our data suggest that TRF can work through the clock if it is present, or reprogram pathways, when it is absent, to provide 24-hour timing in colon tissue that ameliorates colitis.

Our data showing that colitis is improved by TRF is broadly consistent with recent publications^26–29^, however, the use of a 12-hour feeding period is longer than the 4-8 hour feeding used in these previous studies. Specifically, Song *et al*.^26^ reported a 6-hour TRF improved DSS-induced colitis and preserved colon structure, but cell proliferation was not affected. Zhang *et al*.^27^ used an 8-hour TRF and noted reduced inflammation and improved barrier function in this same colitis model. Nui *et al*.^28^ used a 4-hour TRF and found protective effects on intestinal inflammation arising spontaneously in the *IL10* mutant (Crohn’s model) strain; Ruple *et al*.^29^ performed an 8-hour TRF in *IL10* mutants and confirmed the same reduced inflammation. Our data confirm that a 12-hour feeding / fasting regimen recapitulates the anti-inflammatory and colon tissue protective effects observed in much shorter 4-8 hour feeding regimens. A 12-hour TRF diet is a feasible strategy to translate into a clinical setting. The food consumed over a longer feeding window enables individuals to take in the same number of calories (Supplementary Fig 1), thereby avoiding the confounding and potentially negative effects of nutrient restriction.

Colitis is a disease where inflammatory damage to the colon disrupts cellular architecture, that is repaired by the coordinated activities of the epithelium and underlying stroma^57^. In this context, recent studies have shown clock disruption worsens both inflammation and subsequent cellular homeostatic regenerative responses. For instance, we and others have shown that *Bmal1* loss of function severely worsens colitis and prevents recovery^18,28,58,59^. Similarly, the clock components, *Per1/2*, *Nr1d1*, and *Ror* produce severe colitis phenotypes^15–17^. Our present work reveals that TRF can compensate for loss of circadian clock activity in *Bmal1+/+* mice and impose a 24 rhythmicity on the feeding patterns of *Bmal1-/-* mice that benefits colon health. Given the strong link between loss of circadian clock function and increased colitis, TRF-dependent changes to the transcriptional rhythms explain how conferring temporal organization can reduce colitis (Fig 2-3, 5-7).

Imposing a daily nutrition rhythm might also work through the microbiota to improve colitis outcomes. Thaiss *et al*. first reported that *Per1/2* clock mutant mice, who lose rhythms in their microbiota, can have microbiome and transcriptional rhythms restored by timed feeding^34,49^. Indeed, it is known that TRF is able to partially restore 24-hour gene expression rhythms in the liver of *Cry1/2* mutant and *Bmal1* (liver conditional) mutant mice^21^. This suggests time of feeding regulates processes upstream of the resident tissue clocks in digestive system organs. However, recent publications have proposed that epithelial *Bmal1* clock function is required for microbiome rhythmicity^60^ and that it is needed for the TRF-protective effects during colitis^28^. This suggests tissue clocks are upstream of tissue and microbiome rhythms, and that these are not solely driven by food intake. Future studies will resolve these issues, and we note that our present results are consistent with the circadian system driving a rhythm through feeding. In this study, we detected TRF-imposed changes in specific bacterial genera that have been associated with colitis in either humans or animal models. *Ruminococcus* has been noted to increase under TRF^27,61^, and has been reported as a rhythmic^60,61^ bacteria that is reduced under circadian-disruption^49^. *Lachnospiraceae*, is another rhythmic microbe^35,49,60,61^ whose loss of rhythmicity during colitis can be rescued with a 4-hour TRF^28^. *Faecalibaculum* has also been detected^60^, and *Anaeroplasma*^49^ have also been detected as rhythmic. Our work is consistent with these studies and we report additional reductions in *Anaerostipes*, a *Lachnospiraceae* family member that is associated with colitis^54,55^. Our data are consistent with the notion that TRF can alter microbiome composition, and future studies will determine if these relationships are causal or associative with inflammatory disease.

Previous RNA-seq analyses of the intestine or colon have described rhythmic genes related to our findings. Cell division regulators are rhythmic in the colon^34,62^, and interestingly are Bmal1-independent^62^ but microbiome-dependent^34^. Our finding that these are boosted *via* TRF in clock wildtype mice (Fig 3c, 6b-c), suggests that food consumption drives the microbiome to elicit a transcriptional response. Genes related to oxidative phosphorylation, mitochondrial function, and xenobiotic responses are also rhythmic^25,34,36^ and we detect changes in these pathways under feeding and fasting conditions (Fig 6c, 6e, 7c-d).

Diet is a major factor in modulating the gastrointestinal tract, that can be further harnessed to avoid diseases such as colitis. The epithelium and stromal cells of the intestine receive many of their nutrients directly from the lumen, and are thus exquisitely sensitive to daily feeding cycles^63^. Similarly, daily feeding integrates with circadian clock function in intestinal plasma cells to control 24-hour IgA antibody production^64^. Refeeding after fasting induces strong responses from the intestinal epithelium, driving its proliferation through nutrient sensing mTor signaling^65^, and the production of microbial lactate that activates the TCA cycle^66^. Fasting and refeeding can elicit B-cells to immigrate and migrate back to the intestine, respectively, thereby affecting immune system homeostasis^67^. Taken together, these daily responses can be modulated simply through dietary regimens. For example, fasting-mimicking or calorie restriction diets reduce inflammation and promote regeneration in DSS-colitis mouse models^68,69^; this likely contributes to the caloric restriction and colitis improvement we observed in the *Bmal1-/-* mutant (Fig 4). We show here that a 12-hour TRF strategy is sufficient to reduce colitis itself. Future studies will resolve how TRF affects downstream and long-term tissue responses, and whether these can be further optimized using specific nutrition or TRF strategies to prevent IBD.

## Methods

### Animal Housing and Breeding

*Bmal1-/+* mice (Jackson Laboratory, *B6.129-Bmal1tm1Bra/J*, strain # 009100) were housed and bred at the University of Windsor Central Animal Care Facility. *Bmal1+/+* (wild-type, WT) and *Bmal1-/-* (knockout, KO) littermates were co-housed under a 12 hour : 12 hour light-dark cycle (lights on at ZT0; 250 lux) with *ad libitum* access to standard chow (PicoLab-Diet, Cat. No. 5053) and water. All procedures complied with institutional guidelines and were approved by the University of Windsor Animal Care Committee (AUPP 23-17).

At 12-15 weeks of age, mice were individually housed and randomly assigned to either *ad libitum* feeding (AL; 24h food access) or time-restricted feeding (TRF; 12hr nocturnal food access). After 2 weeks, half of the animals were collected and the remaining animals received 4% dextran sulfate sodium (DSS; MW: 40 KDa, Boc Sciences, Cat No. 9011-18- 1) in drinking water for 6 days. Colitis severity was assessed daily during DSS exposure using the Disease Activity Index (DAI)^70^.

### Tissue Collection

Tissues were collected before DSS exposure (d0) and after DSS treatment (d6). *Bmal1+/+* mice were collected at six timepoints including ZT0, 4, 8, 12, 16, and 20 (n = 3 per time point), *Bmal1-/-* mice were collected at two timepoints, ZT0 and ZT12 (n = 3 per time point), for both AL and TRF groups. Colons were dissected and length was measured, after which colons were flushed with ice-cold PBS, opened longitudinally, and rolled from proximal to distal end for histological analysis. Samples were fixed in 4% paraformaldehyde in PBS for 2 hours at room temperature, cryoprotected in 30% sucrose in PBS for 48 hours at 4 °C, embedded in cryo-embedding compound, and stored at −80 °C. Cecum samples were flash-frozen for microbiome analysis.

### Histology

Colon sections (10 µm) were stained with hematoxylin and eosin and imaged using Zeiss Axio Scan.Z1 slide scanner microscope. Lesioned regions of the proximal and distal colon were quantified using QuPath software^71^ by measuring the length of amorphous (lacking crypt structure) regions relative to the total length of each colon. Crypt length was measured in 10 crypts per proximal and distal regions using ZEN software.

### Behavior

Locomotor activity and feeding behavior were assessed in individually housed mice using behavioral chambers (Actimetrics). Locomotion and food intake were recorded continuously for 2 weeks under *ad libitum* feeding conditions, after which half of the animals were switched to a time-restricted feeding (12 hours nocturnal food access). Following an additional 2 weeks, animals received 4% DSS in the drinking water for 6 days. Food intake events were recorded using an automated feeder system dispensing dustless precision pellets (315 ± 4 mg; 3.35 kcal/g; BioServ; Cat. No. F0170). Locomotor and feeding data were collected and analyzed using ClockLab software (Actimetrics)^72^.

To minimize behavioral disruption, DAI scoring was performed at the end of d6. Cage changes were conducted near light offset (ZT12) on day 0, transferring 20% of the original bedding and nest material to the new cage.

### Microbiome Analysis

16S rRNA sequencing was performed on cecum from WT and KO mice. The V3/V4 region was amplified and sequenced via Illumina Mi Seq Illumina as previously described^73,74^. Amplicons were processed by using the DADA2 pipeline and were compared against the SILVA taxonomic database. Analysis was performed using the online tool MicrobiomeAnalyst developed by the Xia Lab at McGill University (https://www.microbiomeanalyst.ca/). Beta diversity was analysed via the Bray-Curtis index, and alpha diversity was analysed using the Shannon index to account for both species’ richness and evenness. The DESeq2 model was utilised to assess differential abundance. Student’s unpaired t-test, one-way analysis of variance (ANOVA) and permutational multivariate analysis of variance (PERMANOVA) were deployed, where appropriate, to assess pairwise comparisons. P values were adjusted using false discovery rate (FDR), and corrected values of <0.05 were considered statistically significant. Full data are provided in Supplementary Table 6.

### Statistical Analysis

All reported experiments used three biological replicates, unless otherwise specified in the figure legend or main text. All histological analysis were performed blinded. GraphPad Prism 9 was used for performing statistical analyses. All data are presented as mean ± s.e.m. Normality was assessed using the Shapiro–Wilk test, and parametric or nonparametric tests were selected accordingly. Brief statistical details are provided in the figure legends, and full details in Supplementary Table 1.

### RNA Collection & Analysis

Colon tissue collected for histology had a 1mm longitudinal segment excised prior to fixation, these samples were stored at −80 °C in RNA stabilization reagent (RNAprotect, Qiagen, Cat. No. 76106) until needed. Tissue samples were homogenized using a Bullet Blender (Next Advance), and total RNA was isolated with the RNeasy Mini Kit (Qiagen, Cat. No. 74106), followed by lithium chloride precipitation as previously described^75^.

RNA samples were submitted to Novogene (https://www.novogene.com) for library preparation and sequencing. Poly(A)+ mRNA was isolated and reverse-transcribed using random hexamers with dUTP-based strand specificity. Libraries were constructed and sequenced on Illumina platforms. Raw FASTQ files were processed with fastp to remove adapters and low-quality reads, these reads were then aligned to the reference genome using HISAT2 with gene model-guided splice junction mapping. All downstream analyses were restricted to protein-coding genes.

### Gene Expression Analysis

Rhythmic gene expression was analyzed using TMM normalized counts. Rhythmicity was assessed using the Metacycle R package^76^. Transcripts with a JTK Benjamini–Hochberg adjusted Q-value (BH.Q) < 0.1 and amplitude < 0.5 were considered rhythmic. Amplitude and phase estimates were used to compare rhythmic parameters across conditions and to perform phase enrichment analysis.

Differential gene expression analysis was performed using DESeq2 (v. 1.44.0)^77^. in R (4.4.1) using the protein-coding universe. Raw count data were pre-filtering by retaining genes with >10 counts in at least 2 samples per group resulting in the removal of approximately 30% of tested genes and a final set of 15,835 genes. A design formula of ∼ 0 + condition was used to enable multiple contrast. Log2 fold-change shrinkage was applied using the ashr method (PMID: 27756721). Genes with a Benjamini-Hochberg adjusted p-value < 0.05 were considered differentially expressed.

Functional enrichment analysis was performed on differentially expressed genes using the clusterProfiler^78^. Gene Ontology (BP) pathway enrichments were conducted. Enrichment analyses applied Benjamini–Hochberg multiple-testing correction (adjusted p-value < 0.05) and gene set size filters (minGSSize = 3); contrasts containing fewer than 10 genes were not tested. Enrichment results were combined per contrast across databases and visualized using dot plots displaying the top enriched terms after de- duplication of highly overlapping gene sets. Conditions tested for differential expression and/or enrichment are indicated in the figure legends.

## Acknowledgments

We thank the technical staff at the Universities of Windsor and McMaster University for their help and advice. VCA, TI, JM, MY, AT, and PK were funded by the Canada Foundation for Innovation, the Ontario Research Fund, and the Canadian Institutes of Health Research. JG, and WK were funded by the Canadian Institutes of Health Research. To the authors whose research we could not cite due to space limitations, we offer our apologies and thanks.

---

**Supplementary Figure 1.**
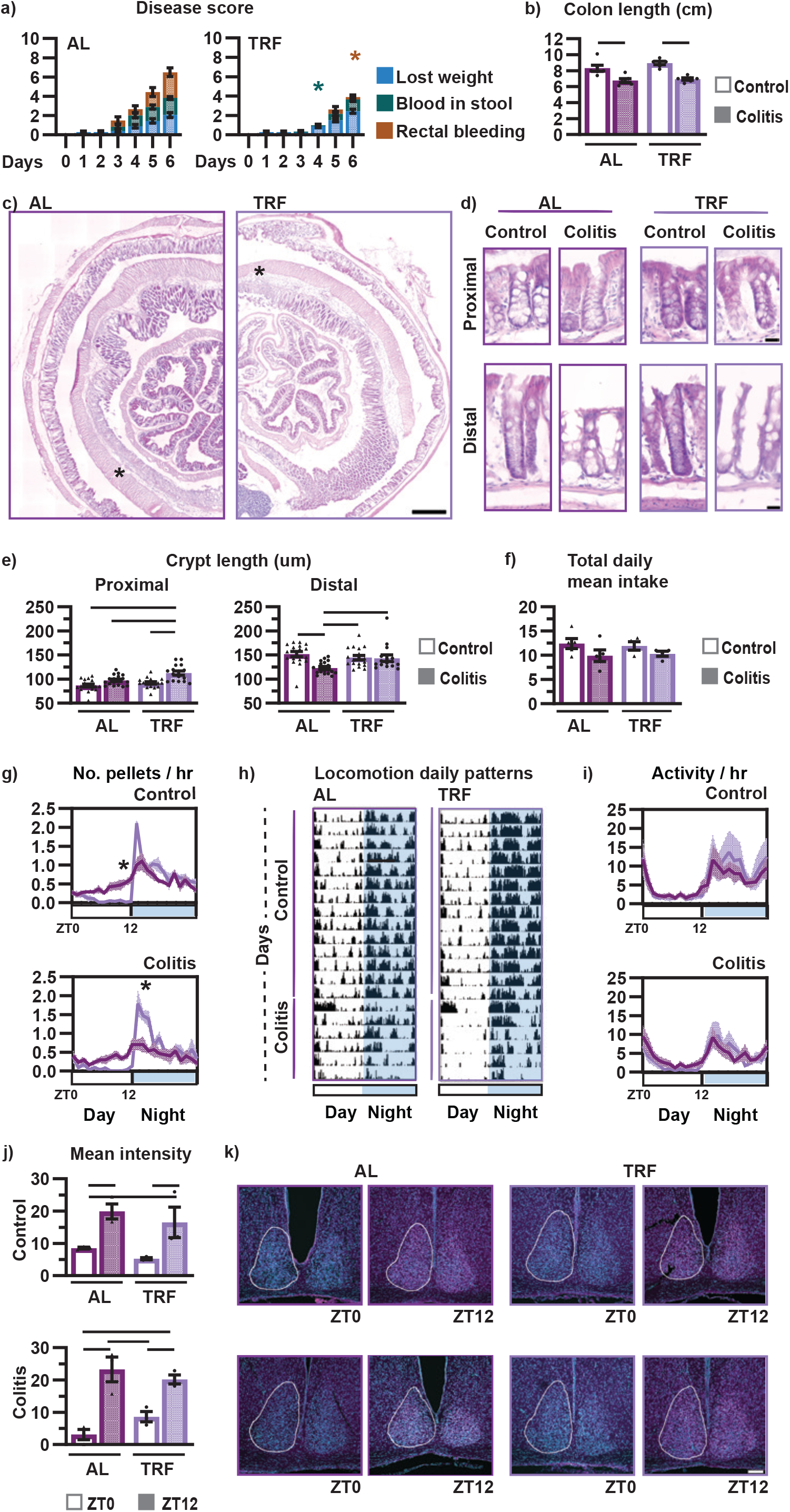
Time-Restricted Feeding (TRF) attenuates colitis independent of total food intake and behaviour. **a)** Differences in disease score between AL and TRF by specific symptoms: lost weight (blue), blood in stool (green), rectal bleeding (brown). Two-way ANOVA repeated measurements, Sidak’s multiple comparisons test. **b)** Colon length decreases during colitis in both AL and TRF conditions. n=6 by group, two-way ANOVA, Tukey’s multiple comparison test (AL, * p=0.0023; TRF, *p= 0.0001). **c)** Representative Hematoxylin and Eosin-stained images showing the distribution of lesion areas across the length of the colon. **d-g)** *Bmal1+/+* mice under TRF show longer crypts during colitis. **d)** Representative H&E from proximal crypts in AL and TRF conditions. **e)** Proximal crypt length is significantly longer in colitis TRF group compared with control AL (p<0.0001), colitis AL (p= 0.0010), and control TRF (p<0.0001, two-way ANOVA & Tukey’s comparison test). **f)** Representative H&E from distal crypts in AL and TRF conditions. **g).** Colitis decreases crypt length compared to controls (p= 0.0002); TRF prevents this decrease in crypt length, there are no significant differences between control TRF and colitis TRF, with colitis TRF showing significantly longer crypts compared to AL colitis (p= 0.0289, two-way ANOVA, Tukey’s comparison test). **h)** Total daily food intake shows no significant differences between conditions (two-way ANOVA). **i)** Daily intake profiles under AL and TRF conditions, before colitis (control, upper panel) and during colitis (lower panel). No significant differences in the daily intake profile were found before (controls) and during colitis across 24-hours. However, we note that under control conditions AL animals increase food intake before the lights go off (ZT10, p=0.0146; ZT11, p=0.0019; ZT12, p<0.0001). During colitis TRF animals eat more at the beginning of the night (ZT12, p<0.0001, ZT13, p=0.0012; ZT14, p=0.0002, two-way ANOVA, Sidak’s multiple comparisons test). **j)** Representative locomotion activity recordings for AL (left) and TRF (right) conditions. X-axis shows hours, ZT0 (onset of the light), ZT12 (offset of the light), Y-axis days across the experiment (Colitis indicated). **k)** No significant differences were found in the locomotor patterns between AL or TRF during control or during colitis conditions. Overall, locomotion decreases during the night during colitis for both AL (ZT16, p=0.0003, ZT17, p=0.0018; ZT18, p=0.0046, ZT22, p=0.0281) and TRF groups (ZT15, p=0.0002, ZT16, p=0.0001; ZT18-19, ZT22-24, p<0.000, two-way ANOVA repeated measurements, Sidak’s multiple comparisons test). Unless otherwise indicated, data are mean ± s.e.m. n=3 by group.

**Supplementary Figure 2.**
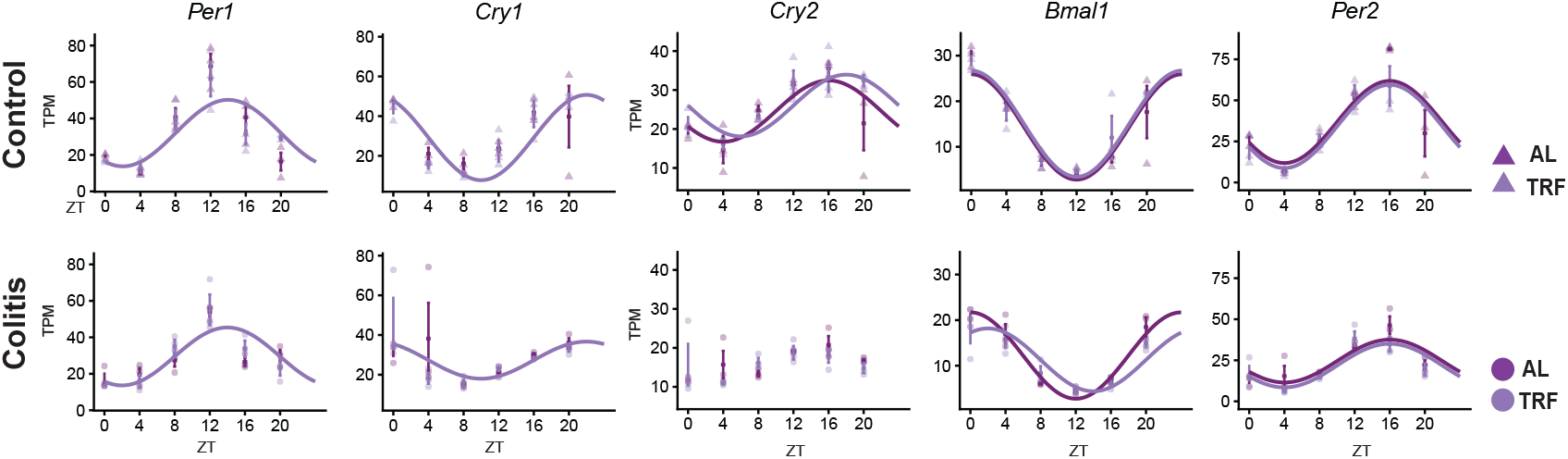
Effects of TRF on the daily rhythms of clock genes in the colon. Daily rhythms in *Per1* and *Cry1* are evident during TRF. During colitis Cry2 results in a loss of rhythms (in both AL and TRF), while *Per2* and *Bmal1* rhythms decrease in amplitude. Graphs show TPM mean ± SEM values across the day. Rhythmic expression was assessed using JTK_CYCLE, and cosine fits are overlaid only for genes classified as rhythmic (BH.Q < 0.1, amplitude > 0.05). Colors represent feeding condition, AL (purple) and TRF(violet), shapes indicate control (triangle) vs colitis (circle).

**Supplementary Figure 3.**
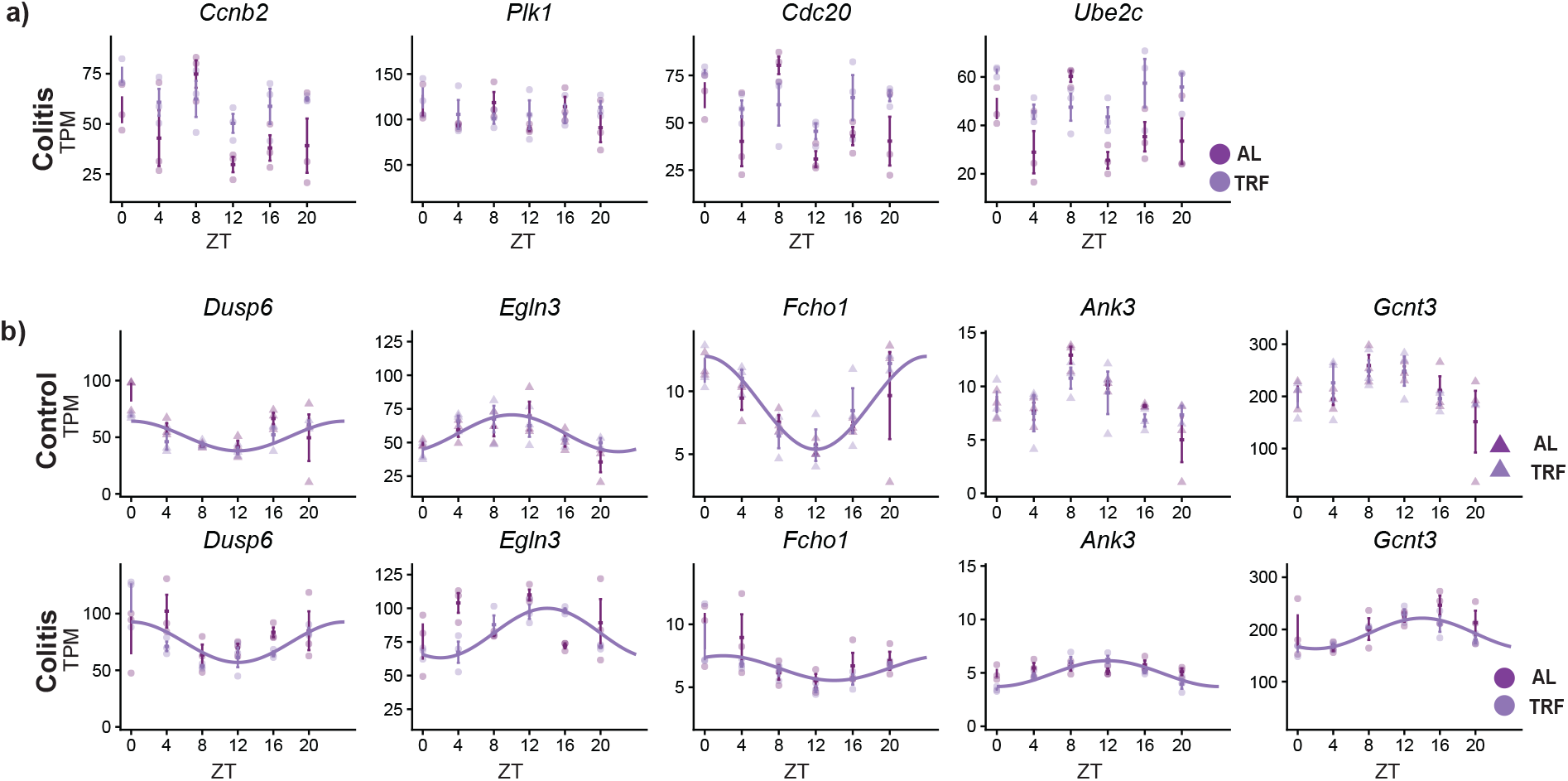
TRF gene expression rhythms. **a)** Examples of genes that lose daily rhythms during colitis. **b)** Examples of genes that have more robust rhythms in colitis under TRF. All graphs show TPM mean ± SEM values across the day. Rhythmic expression was assessed using JTK_CYCLE, and cosine fits are overlaid for genes classified as rhythmic (BH.Q < 0.1, amplitude > 0.05).

**Supplementary Figure 4.**
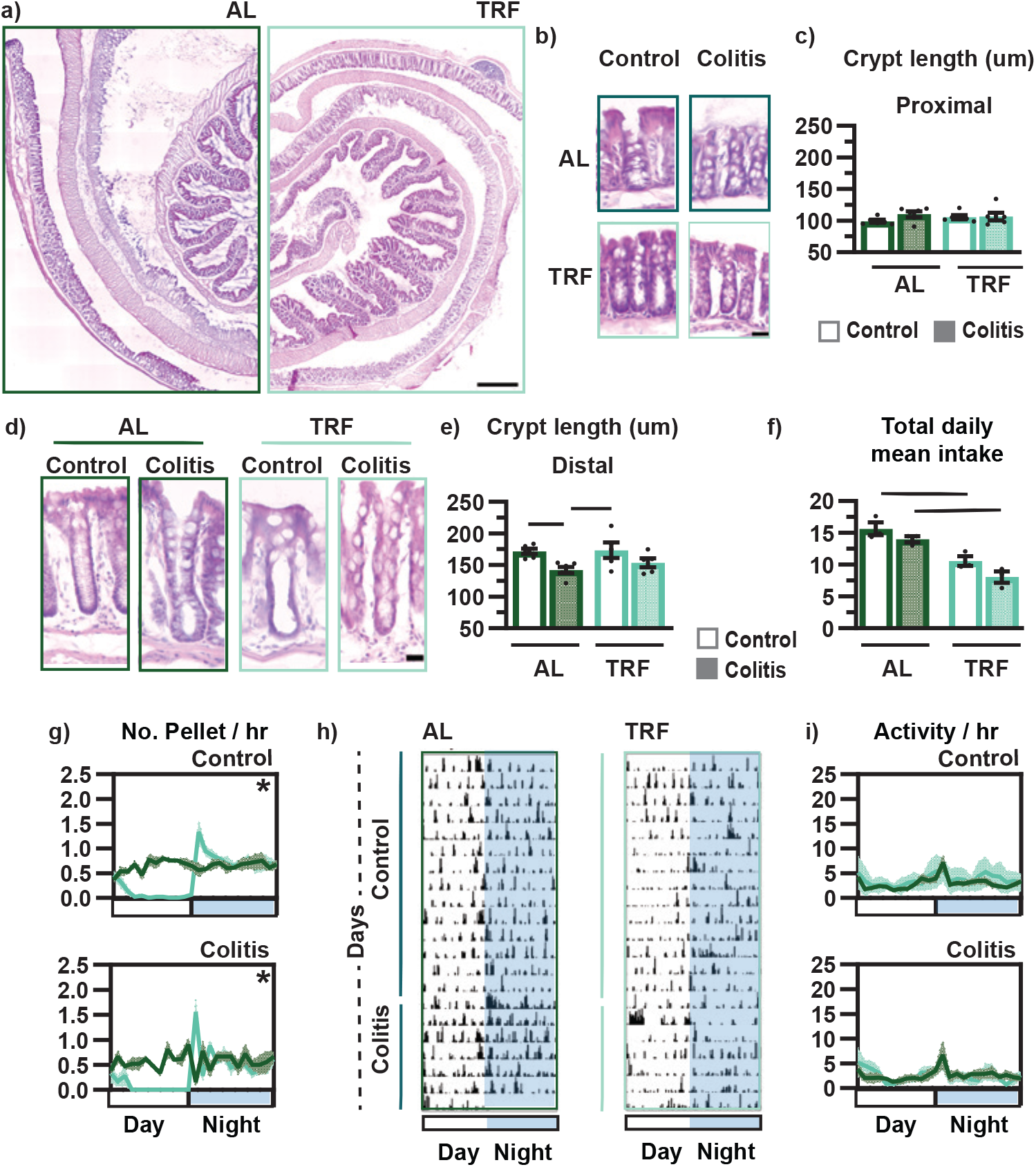
TRF attenuates colitis in Bmal1-/- mutant mice independent of behaviour, but associated with lower total food intake. **a)** Representative H&E showing the distribution of lesion areas across the length of the colon in *Bmal1-/-* mice. **b)** Representative crypts from the proximal colon of *Bmal1-/-* mice in control or colitis conditions, under AL or TRF. **c)** Mean crypt length in the four different conditions, there are no significant differences in the proximal colon (two-way ANOVA). **d-e)** TRF prevents the decrease in crypt length in the distal colon. **d)** Representative H&E from the distal colon across conditions. **e)** Under AL colitis distal colon crypts are significantly shorter (p=0.0265), but TRF prevents this decrease (p=0.0277, two-way ANOVA, Tukey’s comparison test). Unless otherwise indicated, data show mean ± s.e.m., n=6 by group. **f)** *Bmal1-/-* mice under TRF show a decrease in total food intake in control (p=0.0096) and colitis (p=0.0037, two-way ANOVA, Tukey’s comparison test) with respect to mice under AL. **g)** *Bmal1-/-* mice eat continuously across the day under AL conditions. Daily intake profile of control vs. colitis shows how the arrhythmic feeding behaviour of *Bmal1-/-* mice is altered to have a 24-hour feeding rhythm. ***h-i)*** TRF does not alter *Bmal1-/-* arrhythmic locomotor behaviour. **h)** Representative actogram recordings for AL (left) and TRF (right). **i)** No differences were found in the locomotor patterns before and during colitis, between AL and TRF groups (two-way ANOVA). Unless otherwise indicated, data are mean ± s.e.m. n=3 by group.

**Supplementary Figure 5.**
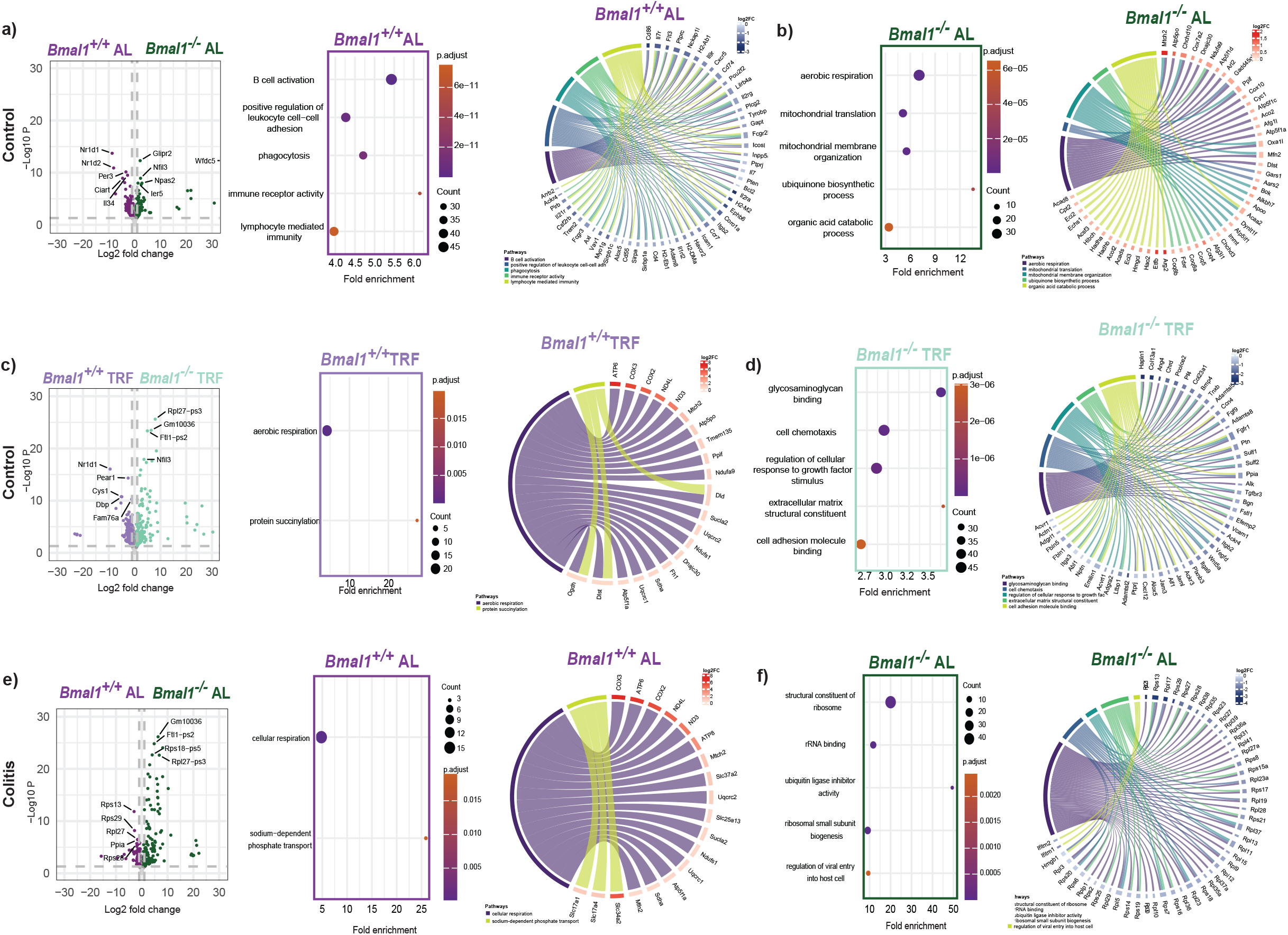
*Bmal1-/-* mice shown metabolic and immune system deficiencies. **a)** Differential gene expression analysis (left) between *Bmal1+/+* AL and *Bmal1-/-* AL under control conditions. Dot plot (middle) shows top 5 enrichment pathways in *Bmal1+/+* AL mice. Chord plot (right) linking differentially expressed genes to enriched pathways, illustrating gene–pathway associations. The *Bmal1+/+* AL colon is enriched for genes regulating immune system activity, these are significantly lower in the *Bmal1-/-* AL colon. **b)** Dot plot (left) of top 5 enrichment pathways in the corresponding *Bmal1-/-* AL mice. Chord plot (right) showing the genes in these pathways as they are linked. The *Bmal1-/-* AL colon is enriched in metabolic processes which are lower in *Bmal1+/+* AL before colitis is induced. **c)** Differential gene expression analysis (left) between *Bmal1+/+* TRF (violet) and *Bmal1-/-* TRF (light green) under control conditions. Dot plot (middle) shows top 5 enrichment pathways in *Bmal1+/+* TRF. Chord plot (right) linking differentially expressed genes to enriched pathways, illustrating gene–pathway associations. The *Bmal1+/+* TRF colon is enriched for genes regulating aerobic respiration (only 2 pathways detected). **d)** Dot plot (left) of top 5 enrichment pathways in the corresponding *Bmal1-/-* TRF mice. Chord plot (right) showing the genes in these pathways as they are linked. The *Bmal1-/-* TRF is enriched in cellular processes including growth factor responses, extracellular matrix, and adhesion which are lower in *Bmal1+/+* TRF before colitis is induced. **e)** Differential gene expression analysis (left) between *Bmal1+/+* AL and *Bmal1- /-* AL during colitis. Dot plot (middle) shows top 5 enrichment pathways in *Bmal1+/+* AL. Chord plot (right) linking differentially expressed genes to enriched pathways, illustrating gene–pathway associations. Following the induction of colitis, the Bmal1+/+ AL colon is enriched in respiration and ion transport (only 2 pathways detected). **f)** Dot plot (left) of top 5 enrichment pathways in the corresponding *Bmal1-/-* AL mice. Chord plot (right) showing the genes in these pathways as they are linked. The *Bmal1-/-* AL colon is enriched in ribosome associated processes which are lower in *Bmal1+/+* AL in the context of colitis.

**Supplementary Figure 6.**
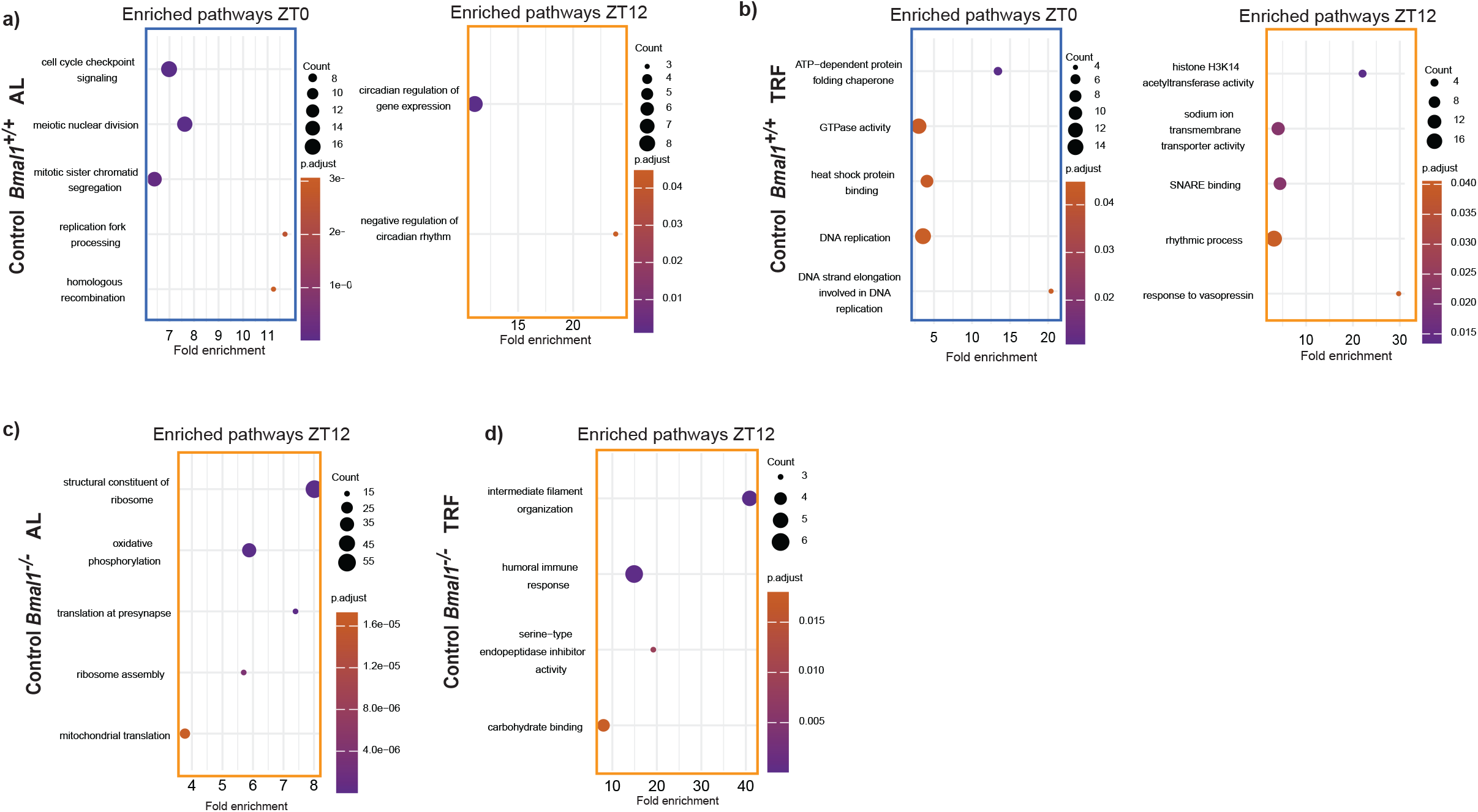
Analysis of enriched Pathways post-feeding *vs*. post-fasting. Dot plots show top 5 enrichment pathways in genes detected by RNA-seq in the different conditions. ZT0 (early morning) following feeding at night, and ZT12 (evening) following fasting all day in the TRF conditions. Enriched pathways correspond to Chord plots shown in Fig 6. **a)** AL control *Bmal1+/+* colon. Cell division pathways are high after feeding and circadian rhythm genes are high after fasting. **b)** TRF control *Bmal1+/+* colon. Cell division and protein production pathways (ATP-dependent proteins folding chaperone, heat shock protein binding) are high after feeding and ion transport pathways (sodium ion transmembrane transport activity, SNARE binding) are high after fasting. **c)** AL control *Bmal1-/-* colon. Only ZT12 shown (no enriched pathways detected at ZT0), where ribosome and metabolic pathways are enriched. **d)** AL control *Bmal1-/-* colon. Only ZT12 shown (no enriched pathways detected at ZT0), where differentiation (intermediate filament organization) and humoral immune responses are enriched.

**Supplementary Figure 7.**
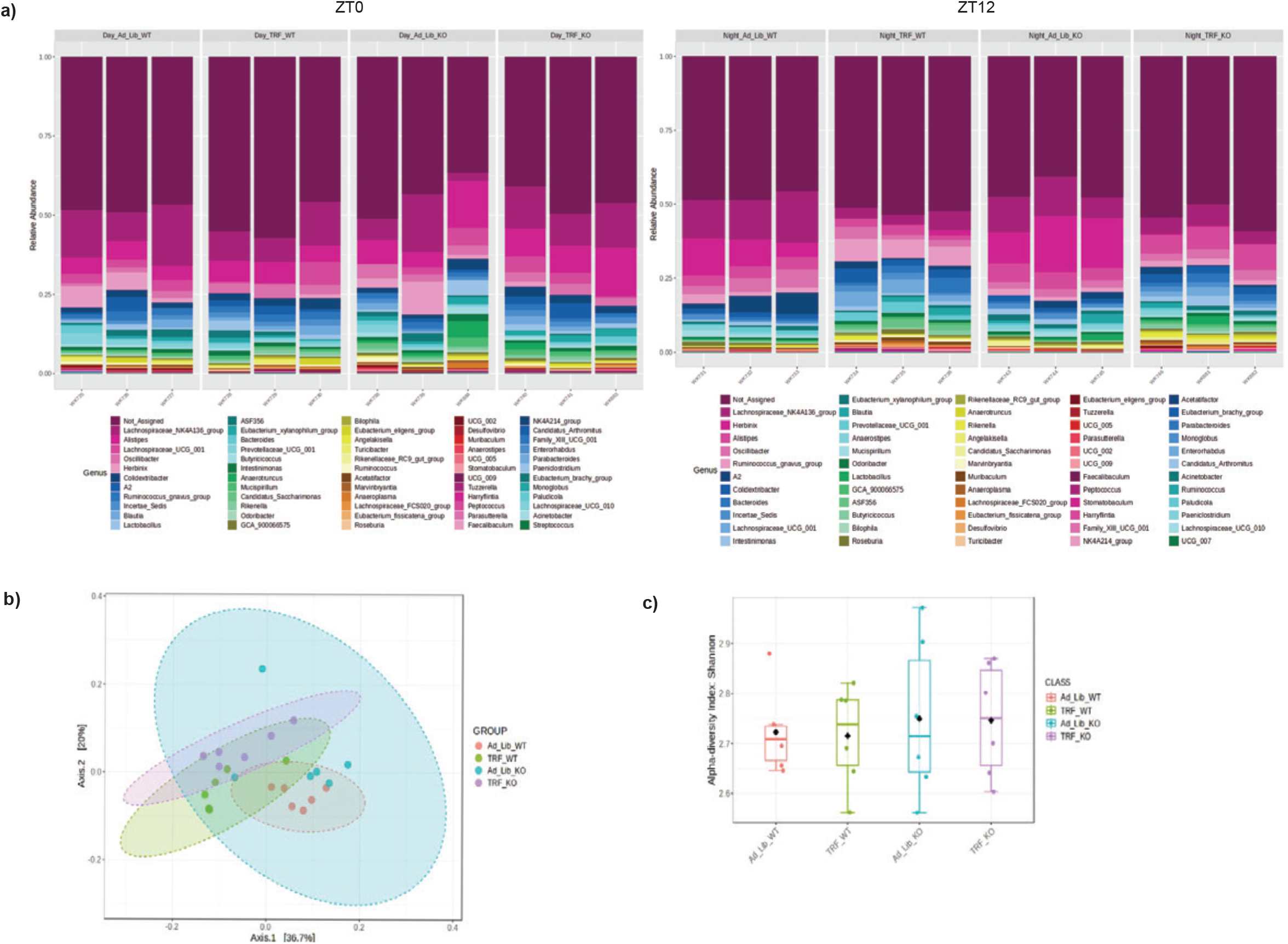
Microbiome analysis. **a)** Microbial relative abundance in *Bmal1+/+* and *Bmal1-/-* mice was assessed by 16s rRNA analysis. All samples were collected prior to colitis (undamaged controls), graphs shown indicate the species detected. **b)** Beta diversity between conditions at the genus level analyzed at ZT0 and ZT12. No significant differences were found. **c)** Similarly, alpha diversity at ZT0 and ZT12 was also not significantly changed by TRF. “Day” refers to ZT0 collections, and “Night” refers to ZT12 collections. “Day_Ad_Lib_WT” refers to *Bmal1+/+* AL, “Day_TRF_WT” refers to *Bmal1+/+* TRF, “Day_Ad_Lib_KO” refers to *Bmal1-/-* AL, “Day_TRF_KO” refers to *Bmal1-/-* TRF.

**Supplementary Figure 8.**
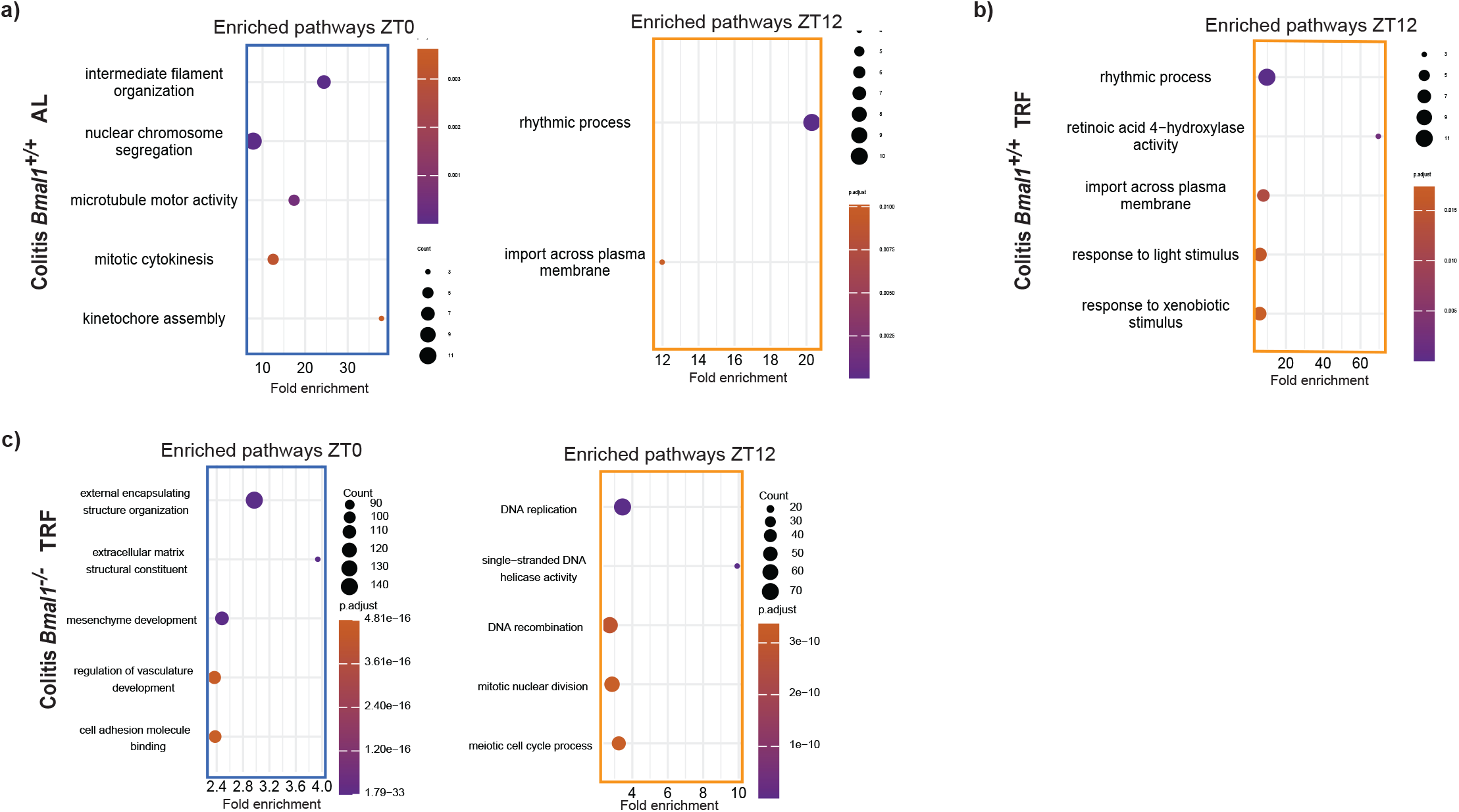
Analysis of enriched pathways in colitis post-feeding *vs*. post-fasting. Dot plots show top 5 enrichment pathways in genes detected by RNA-seq in the different conditions. ZT0 (early morning) following feeding at night, and ZT12 (evening) following fasting all day in the TRF conditions. Enriched pathways correspond to Chord plots shown in Fig 7. **a)** Colitis *Bmal1+/+* AL colon. Similar to control conditions, cell division pathways are high after feeding and circadian rhythm genes are high after fasting. **b)** Colitis *Bmal1+/+* TRF colon shows only enrichment at ZT12 (none detected at ZT0). Rhythmic processes (circadian related genes) and several other processes are significantly enriched at this time. **c)** Colitis *Bmal1-/-* TRF colon (note: no enrichment was observed in the *Bmal1-/-* AL colon during colitis). Extracellular matrix, adhesion, and vascular processes are high at ZT0, while cell division related processes are high at ZT12.

